# Frontal cortex organization supporting audiovisual processing during naturalistic viewing

**DOI:** 10.1101/2025.06.26.661755

**Authors:** Faxin Zhou, Amirhossein Khalilian-Gourtani, Patricia Dugan, Andrew Michalak, Orrin Devinsky, Peter Rozman, Werner Doyle, Daniel Friedman, Adeen Flinker

## Abstract

Our brains dynamically adapt to a multisensory world by orchestrating diverse inputs across sensory streams. This process engages multiple brain regions, but it remains unclear how audiovisual stimuli are represented and evolve over time, especially in naturalistic scenarios. Here, we employed a movie-viewing paradigm to explore this question. We recorded intracranial electrocorticography (iEEG) to measure brain activity in 19 participants watching a short multilingual movie. Using unsupervised clustering and supervised encoding models, we identified a robust modality-specific gradient in the frontal cortex, wherein the ventral division primarily processes auditory information and the dorsal division processes visual inputs. Further, we found that this cortical organization dynamically changed, adapting to different movie contexts. This result potentially reflects flexible audiovisual-resource assignment to construct a coherent percept of the movie. Leveraging behavioral ratings, we found that the frontal cortex is the primary site in this modality assignment process. Together, our findings shed new light on the functional architecture of the frontal cortex underlying flexible multisensory representation and integration in natural contexts.

## Introduction

The brain has a remarkable capacity to process multiple sensory signals simultaneously for a coherent percept [1]. This multisensory processing leverages complementary sensory information to enhance environmental awareness and guide behavior [2]. Neuroimaging and electrophysiological studies have implicated the frontal cortex as a key region in this process [3–5]. Increased activation in the frontal regions occurs when congruent multisensory information is presented [6–8] and when sensory inputs are successfully integrated [9]. Additionally, the frontal lobe also selectively represents relevant information originating from perceptual cortical regions [7, 10].

These findings are primarily derived from highly controlled, task-based experiments that contrast multi- and unisensory conditions. There remains a paucity of research addressing how the brain dynamically organizes multiple sensory inputs in natural settings. To this end, movies can serve as ideal stimuli to study multisensory processing. Movies contain abundant audiovisual information, and observers need to dynamically allocate resources for different modalities to optimize perceptual understanding. Further, naturalistic materials are ecologically valid [11] and potentially generalizable outside the laboratory [12].

Only a limited number of functional Magnetic Resonance Imaging (fMRI) studies have directly investigated audiovisual processing during naturalistic movie viewing. For instance, one employed deep learning (DL)-based encoding models to quantify cortical representations of auditory and visual features and found that audiovisual models provided better fits for frontal cortex versus unimodal models [13]. A related study implicated frontal regions in watching audiovisual movies, as well as in audio-only and visual-only controls, indicating that the frontal cortex represents cross-modal information [14]. These studies, however, did not explicitly differentiate audio and visual representations in the frontal cortex. Moreover, due to the relatively low temporal resolution of fMRI, these studies could not assess moment-to-moment temporal changes in audiovisual processing. Additionally, these studies primarily identified static neural representations, without focusing on how audiovisual representations change throughout the movie. As a result, it remains unclear how the frontal cortex processes auditory and visual information (i.e., in a modality-specific or general manner) and how this functional organization evolves temporally.

To bridge these gaps, we examined neural responses in a cohort of neurosurgical patients using intracranial electroencephalography (iEEG) while they watched a multilingual naturalistic movie. Intracranial EEG directly records from the brain with excellent spatiotemporal resolution (millimeters and milliseconds) and high signal-to-noise ratio (SNR) [15]. Applying functional clustering and encoding models to these recordings, we identified a ventral-to-dorsal gradient in the lateral frontal cortex, associated with auditory (ventral) and visual (dorsal) representations. Further, this modality-specific frontal pattern varied with changes in language context, indicating flexible modality assignment of neural resources. Additional analyses showed that the neural responses related to the audiovisual assignment were predominantly located in the frontal cortex. Overall, these results highlight a dynamic functional organization in the lateral frontal cortex, advancing our understanding of natural audiovisual processing. Our work contributes to the development of neurobiologically plausible models for flexible multisensory processing in real-world scenarios.

## Results

We measured neural activity during free viewing of a movie across 19 patients, including 2688 electrode contacts with wide coverage across the brain (Fig. S1A). The movie contained four different stories with a common theme, alternating between English and other languages (Greek, German, and French; see Methods: Task and procedure). All participants were native English speakers with no prior knowledge of any of the foreign languages in the movie. In this study, we focused on the high-gamma (70 to 150 Hz) broadband field potentials, which are robust at the single-trial level [16], reflect local neuronal firing [17–19], and exhibit strong correlation with the fMRI blood-oxygenation level-dependent (BOLD) response [17, 20].

### Unsupervised clustering reveals modality-specific neural responses

We were first interested in understanding the neural responses to diverse auditory and visual scenes. To this end, we divided the movie into four conditions, each representing a distinct audiovisual scenario (Fig. 1A): (1) English (EN) condition, in which dialogue is spoken in English; (2) Foreign Language (FL) condition, in which characters communicate in non-English languages such as French or German (English subtitles are provided for movie comprehension); (3) Other Sound (OS) condition, in which non-speech sounds co-occur with the visual scene; and (4) Silent (SI) condition, in which visual scenes are presented without audio.

By analyzing neural signals within a one-second post-onset epoch for each condition (onsets were defined as the beginning of the sound in EN, FL, OS or the visual scene in SI, see Methods: Active electrodes selection), we observed various neural characteristics related to audiovisual processing in different regions. For instance, contexts including auditory information (i.e., the EN, FL, and OS conditions) elicited early responses (around 200 ms) in an example electrode from superior temporal gyrus (STG) (Fig. 1C, leftmost panel). Similarly, an electrode in primary visual cortex (V1) exhibited strong activation in the Foreign Language condition in an early epoch (Fig. 1C, the middle left panel). Additionally, frontal electrodes demonstrated delayed responses exclusively for language conditions (after 400 ms), while signals in non-language conditions remained stable (Fig. 1C, the right two panels). The electrodes with significant responses were predominantly distributed in the superior temporal gyrus (STG), occipital regions, frontal cortex, and inferior parietal lobe (IPL; Fig. 1B, S1C, and S8C; permutation test, *p <* 0.05, FDR corrected; see Methods: Active electrodes selection).

To characterize the prototypical patterns of these diverse neural signals, we applied a non-negative matrix factorization (NMF) analysis to the significant electrodes across the four conditions (Fig. S3A, see Methods: Functional clustering analysis) [21–23]. This clustering approach, combined with a silhouette test, identified two distinct clusters (Fig. S3B). The first cluster exhibited higher weights for electrodes in the STG and ventrolateral prefrontal cortex (vlPFC), while the second cluster contained the occipital areas, dorsolateral prefrontal cortex (dlPFC), and pericentral gyrus (Fig. 2A). Based on anatomical structures and their functional responses, we assigned the two clusters as the auditory cluster (red) and the visual cluster (blue), respectively. Furthermore, we calculated the weighted average of neural responses for both clusters (Fig. 2B) and observed that the language conditions generally showed higher amplitudes than non-language conditions. By computing the weighted activity according to distinct anatomical regions (Fig. 2C; Fig. S1B), we found that the auditory and visual clusters showed earlier peaks in the STG and occipital regions respectively (STG electrodes: 365 ± 242 ms; occipital electrodes: 389 ± 221 ms; computed based on the language conditions). In addition, the two clusters showed later responses in the frontal areas (auditory cluster = 657±237 ms; visual cluster peak timing: 608±278 ms; computed based on the language conditions). Notably, the frontal electrodes appeared to be spatially organized according to their respective modalities, wherein the vlPFC electrodes were primarily assigned to the auditory cluster, and the dlPFC electrodes were predominantly assigned to the visual cluster (Fig. 2A). These results indicated a modality-specific cortical activation map during movie viewing.

**Fig. 1:**
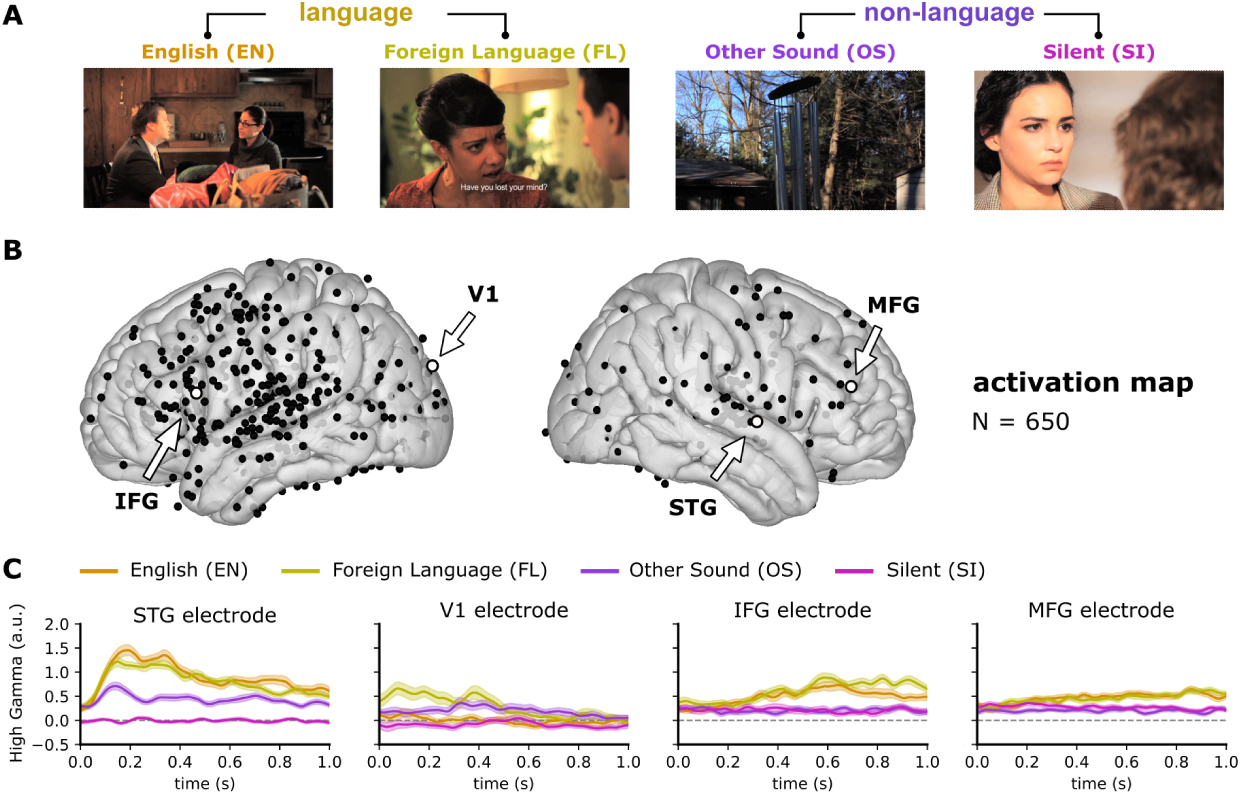
Neural responses during movie viewing. (A) Four conditions identified from the movie: (1) English (EN) condition: dialogue is in English; (2) Foreign Language (FL) condition: communications in other languages such as French or German (English subtitles are provided for movie comprehension); (3) Other Sound (OS) condition: non-speech sounds co-occur with the visual scenes; (4) Silent (SI) condition: visual scenes are presented without audio. (B) All active electrodes survived the permutation test (Fig. S2) across the four conditions (Fig. S1C). (C) Averaged signals across the four conditions for the representative electrodes marked in (B). The shade areas around the curves represent the standard error of the mean (SEM) across trials.

**Fig. 2:**
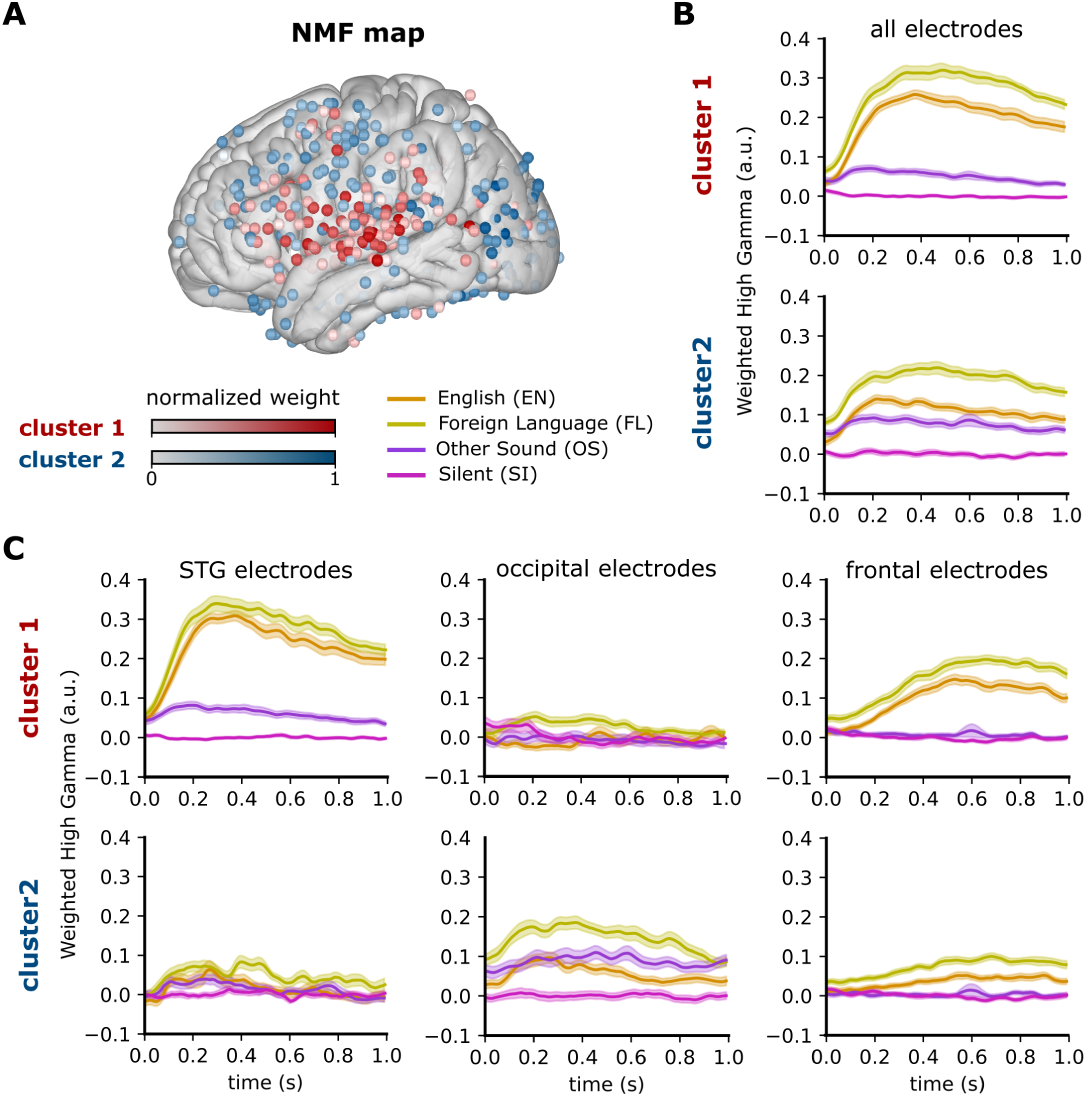
Functional clustering of the active responses. (A) Spatial map showing two identified clusters with the non-negative matrix factorization (NMF) approach. For both clusters, the weights were normalized to the range 0-1, with higher values indicating greater contributions to each cluster. (B) Weighted average signals of the two clusters across all electrodes. (C) Weighted average signals across the electrodes in three specific ROIs (Fig. S1B): STG (left panel), occipital regions (middle panel), and frontal regions (right panel). Shaded areas around the curves represent the SEM.

### Audiovisual encoding exhibits modality-specific representations

We aimed to determine whether the spatial distributions identified by the functional clusters reflect how distinct auditory and visual information is processed during free viewing. To this end, we employed the encoding models to examine how audiovisual information is represented in the brain [24]. First, we utilized a series of computational models to extract low- and high-level features from auditory and visual streams (Fig. 3A, see Methods: Audiovisual feature extraction). At the low level, we were interested in capturing fine-grained acoustic representations from audio (i.e., the spectrogram) and spatiotemporal frequencies from visual scenes (i.e., motion energy Gabor filters [25]). At the high level, by tapping into the significant improvements of transformer architecture in complicated downstream tasks [26, 27], we extracted the vectorial embeddings from two representative models for auditory (the wav2vec 2.0 model [28]) and visual modalities (the vision transformer (ViT) model [29]). Then, we concatenated the low- and high-level embeddings to ensure a comprehensive feature space.

We then used the multivariate temporal response function (mTRF) model to quantify neural encoding [30]. Briefly, the mTRF is a linear model that incorporates time lags (*τ*) between features (*X*) and neural activity (*y*) (Fig. 3B), which has been widely used in neural representational analysis of acoustic, phonological, and linguistic features in connected speech [23, 31, 32] as well as semantic novelty in naturalistic movie segments [33]. To prevent model overfitting and facilitate computational efficiency, we used principal component analysis (PCA) to reduce the dimensions of the audio and visual features (Fig. S4A). We conducted the encoding analysis on all electrodes and evaluated model performance based on the Pearson correlation (*r*) between predicted and actual neural signals in the withheld test set under a 4-fold cross-validation procedure (see Methods: Encoding modeling procedure). Auditory and visual mTRF models were trained separately.

Spatially, significant auditory electrodes were primarily located in STG, vlPFC, and middle precentral gyrus (midPrCG) (Fig. 3C and S5A), while significant visual electrodes were largely distributed in occipital regions, dlPFC, STG, and IPL (Fig. 3D and S5B). The proportion of electrodes tuned to both modalities was relatively low (14.483%) and these electrodes were primarily located in the middle STG, with some scattered across the frontal regions (Fig. S4B). Notably, the anatomical distributions identified by auditory mTRF models replicated our previous NMF analysis of the auditory cluster (Chi-square test, *χ*^2^(9) = 8.921, *p* = 0.445; Fig. S8B, D). Similarly, the spatial patterns identified by the visual mTRF models were consistent with those of the visual cluster, showing no significant difference (Chi-square test, *χ*^2^(9) = 14.337, *p* = 0.111).

Temporally, based on our functional clustering results showing that frontal regions are recruited later than perceptual cortices (e.g., STG and occipital region), we observed similar temporal patterns by examining the lags (*τ*) of the peak weights in the mTRF filters (Fig. S4C; see Methods: Timing analysis of encoding models): the STG (auditory: 194 ± 28 ms) and occipital regions (visual: 224 ± 58 ms) peaked at relatively earlier stages, while the frontal electrodes exhibited later peak latencies (auditory: 396 ± 63 ms; visual: 387 ± 68 ms; Fig. 3E). Moreover, we found no significant differences between modalities in either the lower-level regions (permutation test, *p* = 0.512, FDR corrected) or the frontal regions (permutation test, *p* = 0.745, FDR corrected), but observed a significant difference between them (permutation test, *p <* 0.001, FDR corrected; Fig. 3E). These results collectively suggest distinct encoding of auditory and visual information across multiple timescales.

### A ventral-dorsal gradient in lateral frontal cortex shows audiovisual selectivity

Both the NMF and mTRF results suggest that the ventral and dorsal areas of the frontal cortex are selective for auditory and visual information, respectively. To quantify this relationship, we constructed an index to capture the audiovisual (AV) tuning strength of the significant electrodes in the frontal region. This AV index is computed for each significant electrode by taking the *r* value difference between the visual and auditory models, divided by their sum (see Methods: Frontal ventral-dorsal gradient analysis). Therefore, an AV index of 1 indicates that the electrode is completely visually tuned, while an index of -1 indicates complete auditory tuning. To further delineate the ventral-dorsal gradient, we constructed a frontal polar coordinate system with the intersection of the precentral sulcus and the Sylvian fissure as the origin (MNI coordinate: [-55, 15, -8]). The smaller values of the radius (d, in mm) indicate more ventral positions, while the larger values correspond to more dorsal areas (Fig. 4A). We found a significant linear relationship between the radius and the AV index (*r*(63) = 0.447, *p <* 0.001), quantitatively validating an audiovisual gradient in the frontal cortex (Fig. 4B). Further, we examined what type of audiovisual information drives this frontal gradient. Our feature construction procedure allows us to dissociate low-level perceptual effects (e.g., spectrogram and Gabor features) from high-level representations (e.g., wav2vec 2.0 and ViT features). To assess the contribution of each feature space to neural encoding performance, we computed the reduction in *r*-values when either low- or high-level features were removed from the model (see Methods: Encoding model partitioning). We only observed pronounced effects in the frontal cortex for the features derived from transformer-based models (one-sample *t*-test, auditory: *t*(21) = 9.097, *p <* 0.001, *d* = 1.940; visual: *t*(18) = 4.789, *p <* 0.001, *d* = 1.099; FDR corrected), but not for the low-level features (one-sample *t*-test; auditory: *t*(21) = 1.267, *p* = 0.219, *d* = 0.270; visual: *t*(18) = 2.074, *p* = 0.063, *d* = 0.476; FDR corrected; Fig. 4C). These findings indicated that the frontal gradient is primarily driven by the high-level information embedded in transformer-based models (see Discussion).

**Fig. 3:**
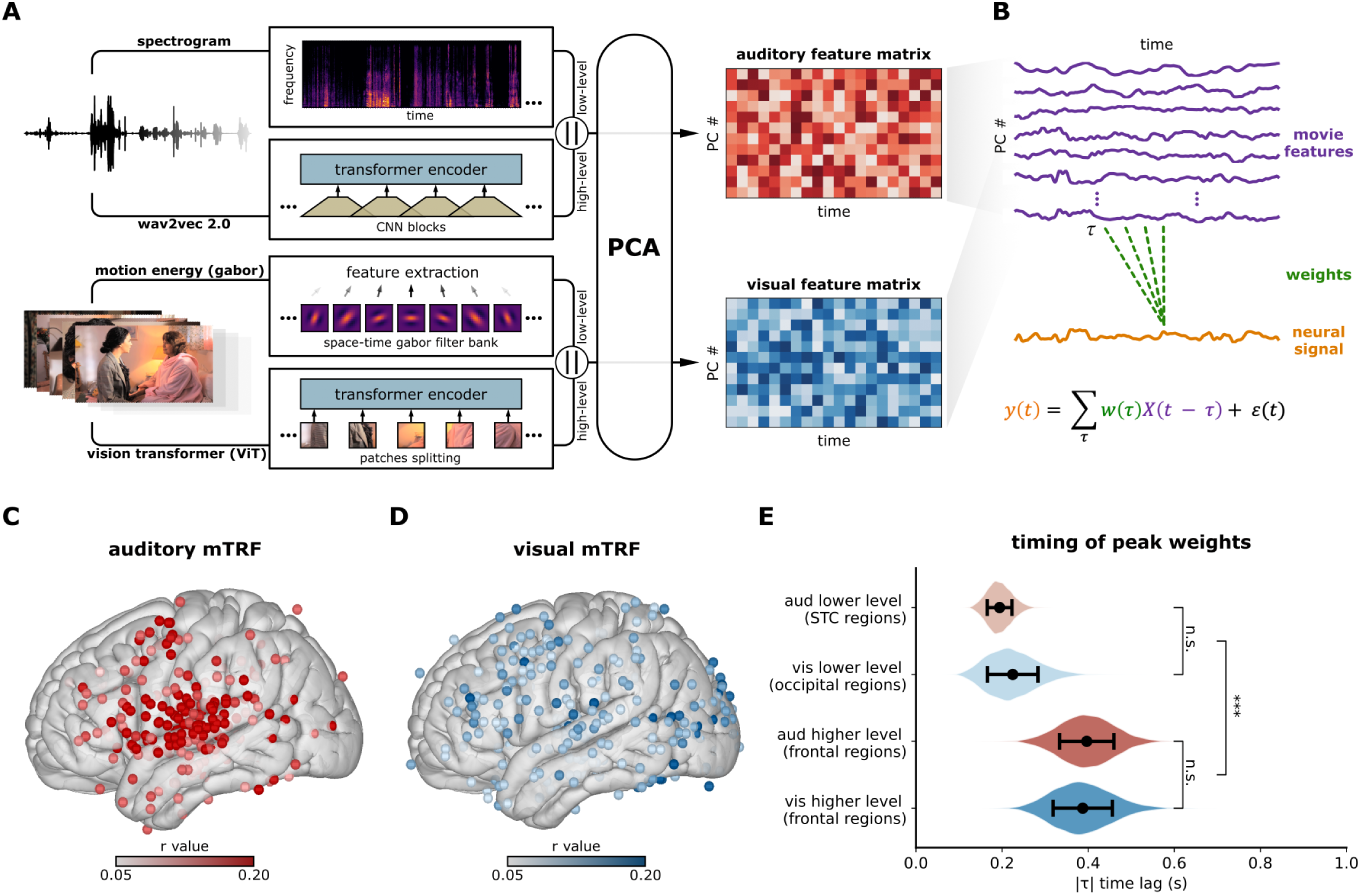
Neural representations of the auditory and visual features in the movie. (A) Pipeline for extracting the audiovisual features, including: 1) low-level auditory (spectrogram) and visual features (motion energy filters); 2) high-level auditory (transformer embeddings from the wav2vec 2.0) and visual features (transformer embeddings from the vision transformer (ViT); see Methods: Audiovisual feature extraction). Low- and high-level features were concatenated (denoted by “||”) before undergoing dimensionality reduction using principal component analysis (PCA). (B) Overview of the multivariate temporal response function (mTRF). The processed movie features (*X*, purple) were used to predict the neural responses (*y*, oranges) with a set of weights (*w*, green) across various time lags (*τ*). (C, D) The correlation brain map of the auditory and visual mTRF models, showing only significant electrodes. (E) The peak timing for all the electrodes in four anatomical region-of-interests (ROIs). The distributions are plotted based on a bootstrap procedure, and the significance test between different regions was conducted with a permutation test (see Methods: Timing analysis of encoding models). The error bars represent the standard deviation (SD). Significance levels are set as *p <* 0.001 (***), *p <* 0.01 (**), *p <* 0.05 (*), and *p* ≥ 0.05 (n.s.).

Additionally, we conducted a control analysis to ensure that the frontal gradient was not biased by the semantic or linguistic components of the subtitles in the Foreign Language condition. By extracting the BERT embeddings of the subtitles and applying a model partitioning procedure (see Methods: Audiovisual feature extraction and Encoding model partitioning), we found that adding the subtitle embeddings neither significantly increased *r*-values in the frontal cortex (Mann-Whitney *U* -Test, *U* (72) = 763, *p* = 0.399, *r* = −0.115; FDR corrected; Fig. 4D-E) nor affected the frontal audiovisual gradient (*r*(47) = 0.594, *p <* 0.001; Fig. 4F-G), indicating that the modality-specific frontal pattern was not confounded by subtitle-related semantic processing.

### A flexible audiovisual-resource assignment supported by behavioral and neural evidence

Next, we were interested in how bimodal information is represented in the brain throughout movie viewing and whether neural patterns vary across different movie context. To this end, we examined the mTRF results for multiple conditions. Although we did not detect frontal representations in the non-language conditions (i.e. Other Sound and Silent; Fig. S5), we observed significant representations changing with language conditions. Specifically, the auditory electrodes were more pronounced in the frontal cortex during the English condition while the visual electrodes were more prominent in the Foreign Language condition (Fig. 5A). Moreover, there was a significant interaction effect between language and modality conditions (two-way ANOVA, modality main effect: *F* (1, 196) = 2.780*, p* = 0.097*, η*^2^ = 0.014; language main effect: *F* (1, 196) = 3.604*, p* = 0.059*, η*^2^ = 0.019; language-by-modality interaction effect: *F* (1, 196) = 55.178, *p <* 0.001, *η*^2^ = 0.220; Fig. 5B), suggesting that audiovisual information is reallocated in the frontal cortex depending on contextual demands. Notably, this pattern was not observed in other areas typically implicated in audiovisual processing, such as the parietal cortex [34, 35], in which auditory encoding performance exceeded visual encoding performance across both the EN and FL conditions, alongside a weaker but significant interaction effect (two-way ANOVA, modality main effect: *F* (1, 122) = 24.340*, p <* 0.001*, η*^2^ = 0.166; language main effect: *F* (1, 122) = 0.448*, p* = 0.505*, η*^2^ = 0.004; language-by-modality interaction effect: *F* (1, 122) = 7.262*, p* = 0.008*, η*^2^ = 0.056; Fig. S7). These results further support the specificity of the frontal cortex as a key site in context switching.

Why did distinct frontal patterns emerge under different language conditions? Past studies suggest that combining multisensory information operates as a weighting process [2, 36–39], indicating that audiovisual processing during movie viewing involves weight-based assignment of different modalities. From this perspective, we would expect the auditory domain to receive greater weights in the English condition and visual cues to be prioritized in the Foreign Language condition, consistent with our findings (Fig. 5A-B). To explicitly test this idea, we conducted a series of movie rating tasks using the Amazon Mechanical Turk (AMT) platform (see Methods: Task and procedure; Fig. 5E). In the task, the entire movie was segmented into 2-3 seconds clips, and participants were asked to evaluate (1) how important the clip is for understanding the entire movie (global context; Fig. 5C); (2) what type of information is more important for understanding the current movie clip (local modality; Fig. 5D). As a result, the point-wise multiplication of the global context and local modality ratings (i.e., the interaction term) could serve as an indicator of modality assignment, quantifying which modality contributes more to overall movie comprehension (global modality; Fig. 5F). Additionally, we performed condition analyses and found that the global context ratings in the language conditions were significantly higher than that in the non-language conditions (two sample *t*-test, *t*(196) = 3.055, *p* = 0.003, *d* = 0.425; Fig. 5G). This result is consistent with our NMF and mTRF findings showing stronger responses for the language conditions (i.e., English and Foreign Language; Fig. 2B and S5). Moreover, our findings for the language conditions also matched our prediction, where the English condition showed significant tuning to auditory information (one sample *t*-test, *t*(43) = −2.562, *p* = 0.014, *d* = 0.386, FDR corrected), while the Foreign Language condition showed greater selectivity for the visual stream (one sample *t*-test, *t*(45) = 5.027, *p <* 0.001, *d* = 0.741, FDR corrected; Fig. 5H).

Further, we reasoned that if this weight-based assignment hypothesis is applicable to audiovisual processing during naturalistic movie viewing, this modality assignment effect should also be detectable at the neural level. To this end, we fitted the mTRF models to the audiovisual assignment variable (i.e., global modality; see Methods: Encoding modeling procedure). Significant electrodes were predominantly located in the lateral frontal cortex and were also sparsely scattered in the anterior temporal lobe (ATL), temporal-parietal junction (TPJ), and other regions (Fig. 6A and S8E). To evaluate the robustness of these findings, we conducted a series of control analyses. First, to rule out the potential confounding effect of attention, we applied the mTRF model to the attentional engagement variable obtained from the engagement rating task (Fig. S9A, B; see Methods: Task and procedure), which revealed a brain pattern distinct from that of the global modality (*r* = 0.015, *p* = 0.431; Fig. S9D). Moreover, the global modality mTRF brain maps trained with and without removing engagement yielded consistent patterns (*r* = 0.621, *p <* 0.001; Fig. S9E, F), suggesting that attentional engagement did not confound our results. Second, we examined whether film cuts could influence the results, as scene transitions might alter the assignment of audiovisual resources. To address this, we computed the first derivative of the global modality and compared it with the film cuts. We defined the film cuts at each abrupt change in movie scenes (i.e., the boundaries; Fig. S9G) [40]. No significant correlations were observed between the film cuts and changes in the global modality across raters (one-sample *t* test against 0: *T* (25) = −1.817, *p* = 0.081, *d* = 0.356; Fig. S9H), indicating that film cuts did not confound the observed effects. Moreover, to exclude potential confounding influences of audiovisual features in the frontal cortex, we performed an additional partitioning procedure (see Methods: Encoding model partitioning) and detected significant effects only for the audiovisual assignment (one sample *t*-test, *t*(33) = 7.968, *p <* 0.001, *d* = 1.366, FDR corrected, Fig. S10A).

Additionally, these electrodes were distinct from those representing audiovisual features. The r-values from the assignment models did not correlate with either auditory (Pearson correlation, *r* = −0.122, *p* = 0.145) or visual r-values (Pearson correlation, *r* = −0.086, *p* = 0.306; Fig. 6B). In contrast to the ventral-to-dorsal frontal gradient observed for the audiovisual features, the electrodes related to audiovisual assignment were more diffusely distributed in the frontal lobe (Fig. 6C). Notably, 73.856% of the electrodes (113 out of 153) were associated with only one specific function, and only four electrodes showed overlap across all three models (2.614%; Fig. 6D). An analysis of the timing (peak lag of modality assignment) showed that the audiovisual assignment in the frontal cortex (297 ± 47 ms) occurred significantly later than the audiovisual representation in the perceptual regions (permutation test, *p* = 0.008, FDR corrected), but earlier than the frontal encoding of audiovisual features (permutation test, *p* = 0.001, FDR corrected; Fig. S10B).

Taken together, our neural and behavioral findings suggest that frontal cortex representations flexibly adapt to shifts in movie plots and linguistic scenarios. Further-more, audiovisual assignment likely relies on distinct frontal neural substrates that complement the representation of audiovisual features.

**Fig. 4:**
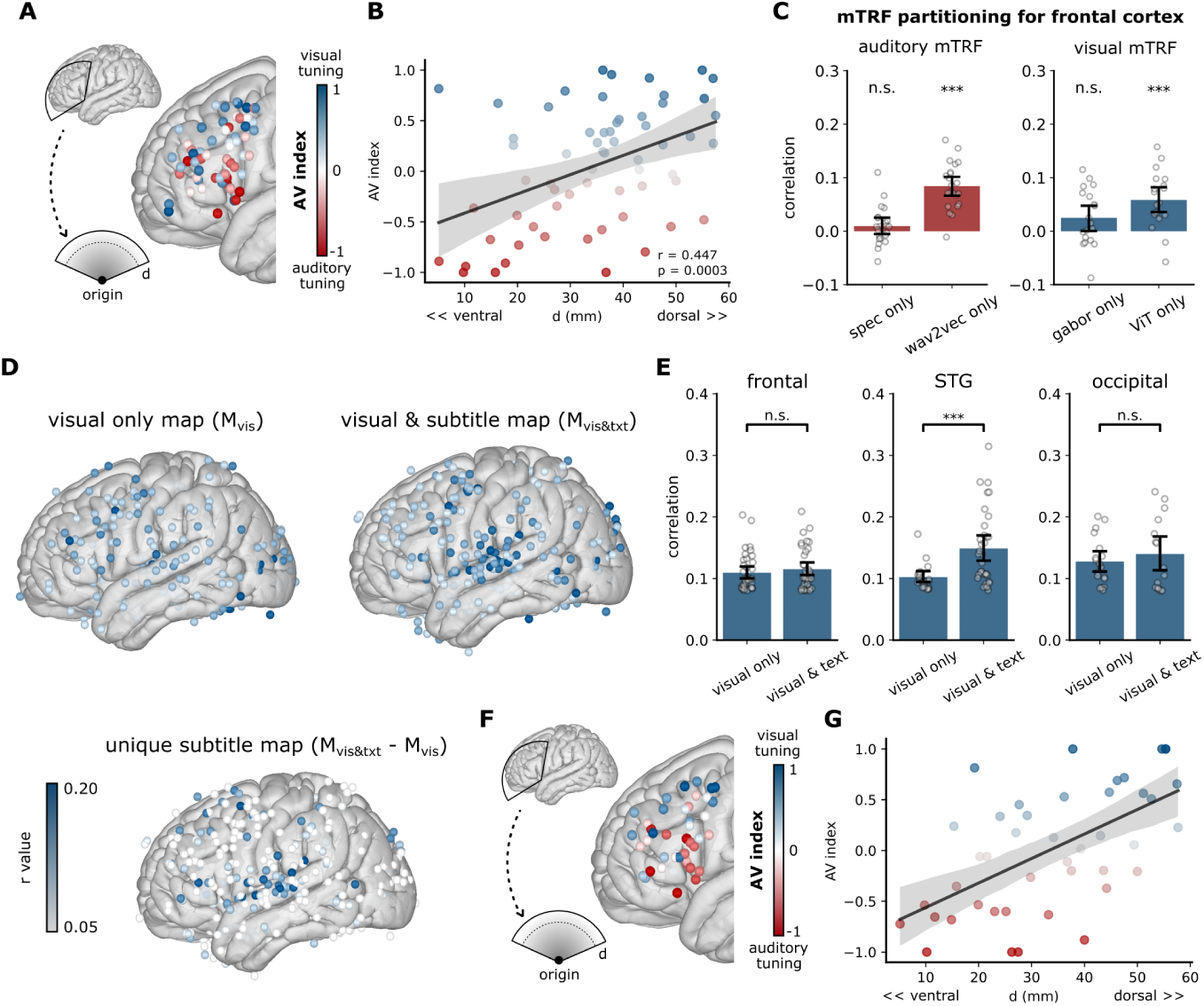
Frontal gradient for audiovisual modalities. (A) The significant electrodes within the frontal sector ROI are represented by audiovisual (AV) index. The sector ROI was defined using a polar coordinate system, with the origin at the intersection of the precentral sulcus and the Sylvian fissure, where the radius (d, mm) quantifies the ventral-to-dorsal gradient. The AV index is computed for each significant electrode by taking the r-value difference between the visual mTRF and auditory mTRF, normalized by their sum (see Methods: Frontal ventral-dorsal gradient anal-ysis). (B) A significant correlation was found between radius and AV index. (C) Results of the encoding model partitioning procedure. We obtained the unique low-level effects (auditory: spectrogram; visual: motion energy Gabor filter) by subtracting the high-level effects (auditory: wav2vec 2.0; visual: vision transformer) from the full model (including both the low- and high-level features). Similarly, the unique high-level effects can be obtained by reversing the procedure (see Methods: Encoding model partitioning). (D) The subtitle effect in Foreign Language condition. Significant neural encoding maps for visual features only (*M_vis_*; top left panel) and for both the visual and subtitle features (*M_vis_*_&_*_txt_*; top right panel). The unique subtitle effect map is the difference between the two maps above (*M_vis_*_&_*_txt_* − *M_vis_*; bottom panel). (E) ROI analysis. The x-axis shows model types (*M_vis_* and *M_vis_*_&_*_txt_*), while the y-axis indicates *r*-values for electrodes within each ROI. No difference was observed in the frontal or the occipital regions. However, a significant increase was found in the STG. (F-G) Frontal gradient analysis incorporating the subtitle embeddings in the visual models (same procedure as figure A-B). The error bars in figure C and E represent 95% CI. The gray area around the regression line in figure B and G represents the 95% confidence interval (CI). Significance levels are set as *p <* 0.001 (***), *p <* 0.01 (**), *p <* 0.05 (*), and *p* ≥ 0.05 (n.s.).

**Fig. 5:**
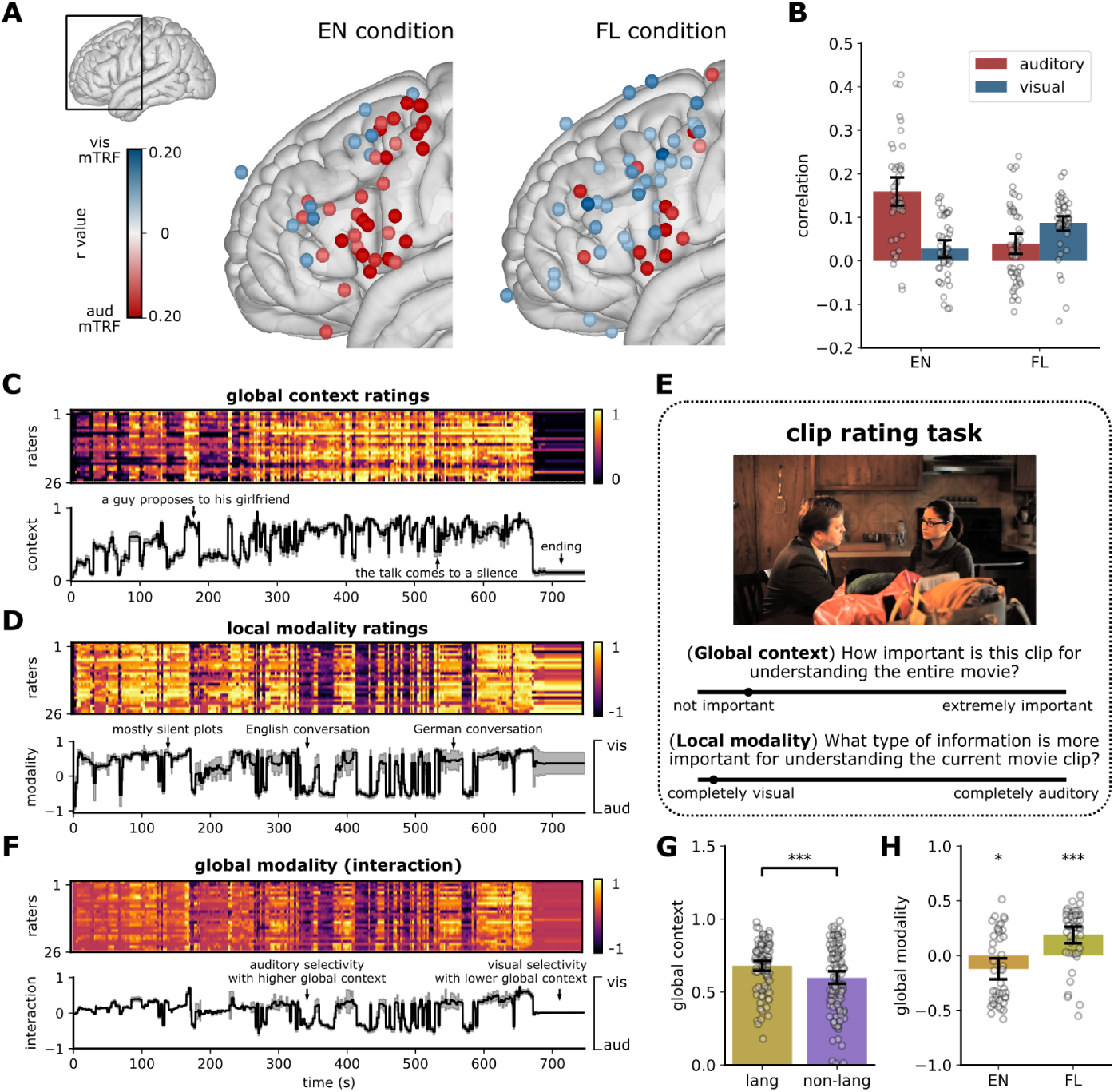
Neural dynamics and the behavior experiment. (A) The mTRF correlation map in the frontal region for English (EN) and Foreign Language (FL) conditions. (B) An interaction effect between the auditory and visual mTRF models in the frontal cortex for the EN and FL conditions. (C, D) The global context and local modality ratings. Upper panel: ratings across all participants over time. Lower panel: the average time series across participants. (E) The pipeline of the clip rating task conducted on the Amazon Mechanical Turk platform. (F) Pointwise multiplication of global context and local modality (the interaction effect). This interaction effect reflects dynamic changes in modality assignment for movie comprehension. (G) The bar plot of the global context ratings for the language (lang) condition, which includes the EN and FL conditions, and the non-language (non-lang) condition, which includes the OS and SI conditions. The dots represent trials in each condition. (H) The bar plot of the global modality for the EN and FL conditions. The shaded areas in figure C, D, and F represent SE; the error bars in figure B, G, and H represent 95% CI. The colors in figure G and H correspond to the four conditions, consistent with the color scheme defined in Fig. 1A. Significance levels are set as *p <* 0.001 (***), *p <* 0.01 (**), *p <* 0.05 (*), and *p* ≥ 0.05 (n.s.).

**Fig. 6:**
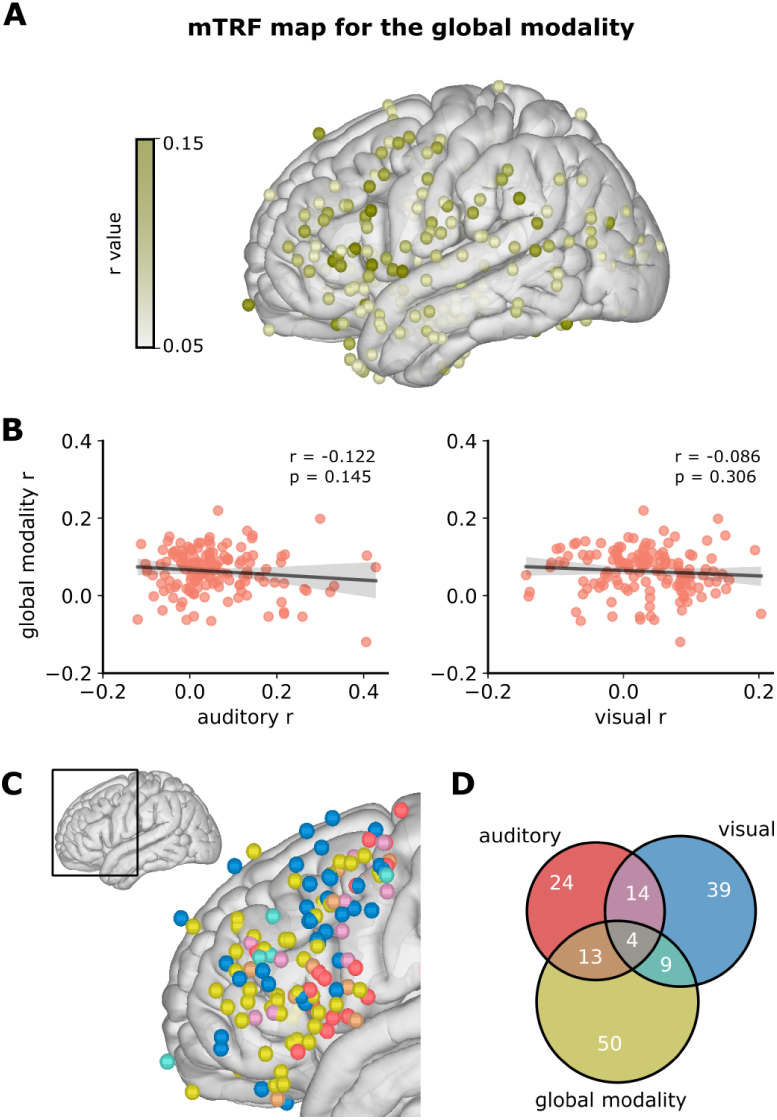
Neural substrates for the audiovisual assignment. (A) The mTRF results for the global modality (i.e., the audiovisual assignment). (B) The correlation analysis between r-values from the assignment mTRF models and r-values from the auditory/visual mTRF models for all significant electrodes in the frontal cortex. The shaded areas around the lines represent the 95% CI. (C) The brain map of significant frontal electrodes across the three mTRF models, rendered with the color code in (D). (D) The Venn diagram of the significant frontal electrodes from the three mTRF models.

## Discussion

The human brain dynamically reconfigures auditory and visual information during free movie viewing. Here, we identified auditory and visual neural responses and representations spanning from the early perceptual areas to the frontal cortex. Notably, the frontal cortex exhibited a modality-specific topography, which shifted depending on the language context. Together with converging behavioral data, we provide evidence that observers dynamically allocate audiovisual resources in response to contextual demands, revealing a flexible and adaptive mechanism for real-world audiovisual processing.

Our results demonstrate a frontal gradient in audiovisual representation during free viewing. Previous studies have shown that the frontal cortex is broadly involved in audiovisual processing during natural movie watching [13, 14] and distinct frontal subregions respond selectively to auditory and visual information in both humans [41–44] and non-human primates [6, 45, 46]. However, these studies primarily focused on specific regions of interest (ROIs) and lacked a comprehensive view of representational organization across the frontal lobe. In contrast, we systematically examined the lateral frontal cortex using both unsupervised functional clustering and supervised encoding models, revealing a robust segregation between auditory and visual modalities. Anatomically, our findings are consistent with the established structural connectivity patterns, wherein the auditory cortices project primarily to the vlPFC (e.g., the IFG) via the arcuate fasciculus (AF) [47, 48] and the visual cortices connect to dlPFC (e.g., the frontal eye field, FEF) through inferior fronto-occipital fasciculus (IFOF) [49]. Functionally, we found that this dorsoventral gradient was predominantly driven by high-level features extracted from transformer-based models and exhibited relatively late peak latencies (around 400 ms; Fig. 4C). Moreover, the sustained frontal activity during the language context (Fig. 2C) suggests its involvement in maintaining recent narrative information over time [50, 51]. These findings align with recent unimodal auditory and visual studies reporting dissociable semantic representations in the frontal cortex [21, 52–56] as well as studies combining structural and functional connectivity showing a dorsoventral frontal gradient corresponding to an auditory-to-visual transition [57]. Extending these findings, our study provides the first direct iEEG evidence for an audiovisual semantic gradient along the dorsoventral axis in frontal regions during naturalistic movie viewing.

Importantly, rather than restricting our analysis to predefined anatomical areas, we constructed a frontal sector coordinate system and identified a modality-specific gradient, transitioning from the ventral region associated with auditory information to the dorsal region associated with visual information. By constructing a continuous ventrodorsal index from our coordinate system, we were able to quantify modality transition trends in a fully data-driven manner without imposing rigid anatomical assumptions. This suggests a spatially continuous coordinate system along the ventral–dorsal axis of the frontal cortex [58–60], reflecting a functional topographic organization that supports audiovisual processing under naturalistic experiences.

In addition, prior perceptual research has shown that observers integrate multimodal cues by assigning weights proportional to their reliability across different task contexts [34, 38, 39, 61, 62]. Under this view, more reliable modalities tend to exert greater influence on unified perception. Our findings suggest a similar “modality assignment strategy” during movie viewing, in which frontal neural resources are dynamically allocated between auditory and visual modalities based on contextual demands. Unlike prior studies that manipulated stimulus reliability by adding noise, movies are carefully crafted and contain minimal artificial distortion [11, 63], with both auditory and visual streams maintaining consistently high fidelity. Therefore, the modality assignment effect we observed is unlikely to stem from sensory reliability *per se*, but instead reflects how different modalities contribute to forming a coherent perceptual object. This aligns with a framework of goal-driven, top-down control over multisensory processing [64]. However, the modality assignment effect could also reflect the influence of attention induced by language intelligibility. To exclude this alternative explanation, we conducted a series of control analyses (Fig. S9A-F), which demonstrated that attentional engagement did not significantly contribute to the observed modality assignment effect. Moreover, we found that the modality assignment effect peaked around 300 ms, consistent with the temporal window associated with multisensory integration [65–67]. The frontal cluster associated with the “modality assignment” also aligns with prior evidence implicating frontal regions in processing novel information during multimodal integration [68–70]. Collectively, our results provide an expansion from previous well-controlled laboratory experiments to naturalistic settings, revealing a goal-oriented modality assignment strategy for audiovisual integration during free viewing of movies.

Several limitations should be noted. First, the constraints of bedside iEEG experiments in the hospital (e.g., variable distance between participants and the laptop) may introduce variability in cortical activity and reduce SNR. Second, the absence of behavioral measures from patients (e.g., eye-tracking recordings and direct assessments of movie comprehension) limited our ability to directly account for attention and eye movements. Third, unbalanced electrode coverage across hemispheres (Fig. S1A) precluded an investigation of the potential lateralization effects during movie viewing. Lastly, vlPFC responses were primarily observed during intelligible speech, raising the possibility that the frontal audiovisual gradient may be influenced by language-related processes rather than purely audiovisual. The present study did not include naturalistic stimuli lacking linguistic content, thus we could not rule out this interpretation, and we believe that the relationship between language and audiovisual processing warrants further investigation.

In summary, our study demonstrates that the lateral frontal cortex serves as a key region for processing audiovisual information under naturalistic experiences. We reveal modality-specific representations in the frontal cortex, with auditory information encoded toward ventral regions and visual information toward dorsal areas. We also show that distinct frontal substrates are largely involved in the modality assignment process. Together, these findings provide evidence for a distinct spatial organization in the frontal cortex supporting flexible multisensory processing of real-world scenarios.

## Methods

### Participants

The first cohort consisted of 19 patients undergoing neurosurgical evaluation for refractory epilepsy (12 females; all right-handed; 32.053 ± 11.297 years old; 12 grid and 7 stereotactic coverages; see Supplementary Table S1). These patients participated in a multilingual movie watching task during their stay in the hospital. Electrode implantation was determined exclusively based on clinical requirements. All participants were fluent in English and had no knowledge of the foreign languages used in the movie. Written informed consent was obtained from all participants either prior to the neurosurgical procedure or upon admission to the hospital unit for evaluation. A second cohort of 26 crowdsourced raters was included in an online movie rating task (11 females; 3 left-handed; 42.154 ± 8.694 years old). Participants self-identified as either native English speakers (23 raters) or proficient English speakers (3 raters), none reported knowledge of the foreign languages used in the movie.

The current study protocol was approved by the NYU Langone Medical Cen-ter Committee on Human Research. All participants were reimbursed for their participation.

### Task and procedure

In the intracranial experiment (**Movie watching task**), patients were asked to silently watch a 12-minute long multilingual movie called Foreign Language Movie (FLM; https://vimeo.com/61040183). This short movie contains four distinct story-lines: 1) a Greek mother reveals her terminal disease to her daughter (Greek and English); 2) two German girls discuss saving an unborn life (German); 3) a married couple’s relationship comes to an end (English); 4) a young couple conversing in French and English about getting married. The stories are intricately interwoven, forming a cohesive and unified story. English subtitles are provided in the foreign language conversations to aid in movie comprehension.

The movie was presented on a 15” laptop placed 0.5–1.0 meters in front of the participants. Audio was delivered through a nearby speaker beside the laptop, with the volume adjusted to a clear and comfortable level for each participant. To synchronize neural activity with the movie, trigger pulses were transmitted to the EEG recording system at the movie onset and at one-minute intervals thereafter. For all included patients, the experiment was not intervened by clinicians or relatives.

The behavioral experiment (**Online movie-rating task**) was conducted using the Amazon Mechanical Turk (AMT) platform and required participants to watch the movie twice. Before the experiment, raters were instructed to “find a quiet place, make sure that your internet has a stable connection and will not be disturbed during the experiment”. During the first viewing, participants watched the entire movie and were asked to provide continuous engagement ratings by adjusting a scale bar from 1 (not engaging) to 99 (extremely engaging) [71]. The scale bar remained visible below the video window throughout the session. If participants did not rate for longer than 5 seconds (i.e., did not move the cursor), the movie would automatically pause, and a flashing reminder would prompt them to rate. Immediately after the first viewing, participants completed a free-recall task to assess comprehension. The instruction for this section was “Please describe what you remember from the movie. Please try to recount events in their original order, if possible, and describe events in writing in as much detail as possible (spend approximately 5-10 minutes). During your description, completeness and detail are more important than the order of events, typos, and grammar.” [72]. Raters whose responses lacked sufficient detail or omitted one or more storylines were excluded from further analysis.

During a second viewing, the movie was presented as a sequence of 2-3 second clips. The segmentation was performed using the FFmpeg toolbox (https://www.ffmpeg.org/). For each clip, the raters were asked to evaluate two aspects using scale bars: 1) how important is this clip for understanding the entire movie from 1 (not important) to 99 (extremely important); 2) what type of information is more important for understanding the current movie clip from 1 (completely visual) to 99 (completely auditory) (Fig. 5E). If unsure of their ratings, the raters could replay the clips before proceeding to the next clip. The resulting clip-specific ratings were concatenated (based on frame counts) to reconstruct continuous movie rating signals (Fig. 5C-D).

### Statistical testing

For both behavioral and neural data, the D’Agostino test was used to determine the normality of the distribution prior to statistical analysis. If data followed a normal distribution, a parametric test was used (e.g., paired *t* -test); otherwise, a non-parametric test was used (e.g., Wilcoxon signed-rank test). In addition, we applied a permutation test to identify active electrodes (e.g., Fig. 1B) and assess significance in the encoding models (e.g., Fig. 3C-D). Detailed procedures are described in the corresponding Methods sections (see Methods: Active electrodes selection and Encoding modeling procedure).

All statistical tests were two-tailed with a significance threshold of *p <* 0.05, unless stated otherwise. The false discovery rate (FDR) correction was applied when multiple comparisons were conducted.

### Data acquisition and preprocessing

During the movie watching task, iEEG signals were recorded using one of two amplifier types (dictated by clinical location during acquisition): (1) NicoletOne amplifier (Natus Neurologics, Middleton, WI), with signals bandpass filtered from 0.16 to 250 Hz and digitized at 512 Hz; (2) Neuroworks Quantum Amplifier (Natus Biomedical, Appleton, WI), with signals recorded at a sampling rate of 2048 Hz, bandpass filtered between 0.01 to 682.67 Hz and then decimated to 512 Hz. For grid electrodes, a two-contact subdural strip facing the skull near the craniotomy site served as the reference electrode, and a similar strip screwed to the skull was used as the instrument ground. For sEEG electrodes, a five-lead subgaleal strip was placed facing the skull and used as both the ground and reference. Electrodes within the seizure onset zone (SOZ; Fig. S1A), or showing epileptiform activity, line noise artifacts, or large amplitude shifts were excluded. The data were then re-referenced using the common average reference (CAR) approach, in which the averaged signal was subtracted from all electrodes for each subject. To extract high-gamma broadband activity (70–150 Hz), we applied a multi-band averaging method. Specifically, we computed the z-scored analytic amplitude across eight logarithmically spaced frequency bands within this range using the Hilbert transform, and then averaged them [21, 73]. This method enhances signal robustness as it takes the mean across multiple frequency bands and mitigates the bias of lower frequencies given the z-score normalization. Data were then corrected relative to the baseline (with silent, black scenes) for each channel.

Electrode localization in individual space was performed by co-registering post-operative brain MRI or CT scans to preoperative MRI scans using a rigid-body transformation. Electrodes were then projected into the MNI space using a nonlinear DARTEL algorithm [74]. Further, anatomical locations of electrodes were determined using automated FreeSurfer segmentation of the preoperative MRI. Visualization of electrode locations on the cortical surface was performed using Mithra [75].

### Active electrodes selection

To identify the active electrodes involved in audiovisual processing during free viewing, we extracted the neural responses in four different audiovisual conditions (Fig. 1A): 1) English (EN) condition, in which dialogue is in English; 2) foreign language (FL) condition, in which characters communicate in other languages like French or German (English subtitles are provided to aid movie comprehension); 3) other sound (OS) condition, in which non-speech sounds are presented with visual scenes; and 4) silent (SI) condition, in which only visual scenes are presented without sound. Specifically, for the conditions with sound (i.e., EN, FL, and OS), we manually annotated the trial onsets and offsets based on the sound waveform and spectrogram using the Praat software (https://www.fon.hum.uva.nl/praat/). The remaining intervals were then labeled as the silent period. To minimize the potential residual activity, we marked the onsets of silent periods 200-500 ms after the preceding sound offsets. Trials shorter than one second were excluded to ensure data stability and robustness. As a result, 44 EN trials, 46 FL trials, 58 OS trials, and 51 SI trials with comparable segment durations were included for analysis (one-way ANOVA: *F* (3, 198) = 1.020, *p* = 0.385, *η*^2^ = 0.061; Fig. S3C).

For each electrode, we aligned the onset of all trials and analyzed the potential changes within a one-second epoch in each condition. An electrode was considered active if its activity across trials was significantly greater than zero (one sample *t* - test) for a duration exceeding a threshold determined by the permutation procedure (Fig. S2). Specifically, in each permutation, we distorted the temporal alignment of the signals using a phase randomization approach [76], then identifying the maximum number of consecutive significant time points (i.e., significantly above zero) using the same analysis. Repeating this procedure 1000 times yielded a null distribution of significant duration. Then, the *p*-value can be derived by calculating the proportion in the null distribution that exceeded the real significant duration. An electrode was considered significant if *p <* 0.05 and survived the false discovery rate (FDR) correction of multiple comparisons (Fig. S2C). The union of electrodes identified as active across the four conditions was used for subsequent analyses.

### Functional clustering analysis

We performed unsupervised non-negative matrix factorization (NMF) to summarize neural underpinnings associated with audiovisual processing [21, 23]. By introducing the non-negativity constraint, NMF is able to effectively derive the parts-based representation [77]. Mathematically, a non-negative data matrix (**D** ∈ R*^m×n^*) can be factorized into two components **W** ∈ R*^m×k^* and **H** ∈ R*^k×n^* by minimizing the error function iteratively (using the Frobenius norm):

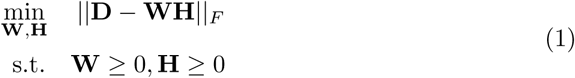

where the term *k* is a hyperparameter representing the number of factorized clusters. In practice, the matrix **D** represents the concatenated neural responses of active electrodes across four conditions (Fig. S3A). We imposed non-negativity in **D** by setting all negative elements to zero, as no significant negative activations were identified with the permutation test (see Methods: Active electrodes selection). **W** is the weighting matrix, representing the extent to which electrodes were assigned to each cluster. **H** is the temporal prototype matrix, demonstrating the typical temporal dynamics for each cluster. To identify the best number of clusters (*k*), we applied the silhouette method to measure the similarity of each sample to its own cluster relative to others [78]. The highest silhouette score, which indicates the optimal clustering assignment, was found when *k* = 2 (Fig. S3B). Further, we observed that the two factorized clusters were largely independent with minimal overlap (Fig. S3D). Therefore, electrodes were assigned to clusters based on their maximum contributions in the weighting matrix **W**, and were normalized (0-1) for visualization purposes (Fig. 2A).

### Audiovisual feature extraction

We employed several computational models to derive generalizable feature spaces for both auditory and visual modalities (Fig. 3A). Specifically, we extracted features at two levels. At the low levels, we captured the spectrogram and motion energy features to represent auditory and visual information, respectively. For the spectrogram, we applied the librosa Python package [79] to extract spectral frequencies ranging from approximately 20 to 8000 Hz, which serves as an effective acoustic representation of auditory perception. For the motion energy features, we utilized a pyramid of non-linear spatiotemporal Gabor filters, which are designed to capture information across multiple positions, orientations, spatiotemporal frequencies, and motion directions [25]. This filter bank spans both small and large receptive field sizes, making it sensitive to both local changes and global motions. The feature extraction procedure was implemented with the pymoten package (https://github.com/gallantlab/pymoten).

At the high levels, we leveraged transformer-based deep-learning models, which have demonstrated significant improvements in complicated downstream tasks [26, 27]. The core mechanism of the transformer architecture is multihead self-attention, an algorithm that calculates context-dependent weights to integrate information across input vectors [80]. In practice, we extracted features from two representative models: wav2vec 2.0 for auditory information [28] and Vision Transformer (ViT) for visual information [29]. To avoid task-specific influences and capture representative features of audiovisual information, we selected vector embeddings from the middle layers, as prior research indicated that these layers produce the most comprehensive features for both the wav2vec 2.0 [53, 81] and the ViT models [82]. In detail, for the auditory domain, we extracted 7th layer embeddings of transformer blocks in wav2vec 2.0-Base model (number of convolution layers = 7, number of transformer layers = 12, hidden size = 768, number of self-attention heads = 8, total parameters = 95 M) [28]. We kept the same sampling rate (16 kHz) and batch size (15.6 seconds) with the pretrain models during feature extraction to minimize potential input format biases. For the visual domain, we extracted the [CLS] embeddings from the 16th layer of the ViT-Huge model (number of transformer layers = 32, hidden size = 1280, number of self-attention heads = 16, total parameters = 632 M) [29]. [CLS] is an extra learnable “token” introduced to the transformer encoder, representing image representations for the classification task.

Furthermore, to control the potential effect of visual subtitles in the Foreign Language condition (Fig. 4D), we extracted semantic embeddings of these sentences utilizing the robustly optimized BERT approach (RoBERTa) model [83]. RoBERTa is a generalized BERT representational model, featuring a multi-layer bidirectional transformer encoder conditioned on both left and right contexts [84]. Compared with the original BERT model, RoBERTa model has a larger architecture, with a bigger batch size and more training data. Here, we applied the RoBERTa-Large model (the number of layers = 24, the hidden size = 1024, the number of self-attention heads = 16, total parameters = 355 M) [83], and selected the features from the penultimate hidden layers, as the final layers tend to be biased by the model training objectives (i.e., masked language model and next sentence prediction). Practically, we represented sentence embeddings by averaging all word vectors within each sentence [85–87].

All feature extraction procedures associated with deep-learning models were performed using the PyTorch (v2.1.0) framework [88] in Python (v3.10).

### Encoding modeling procedure

We utilized encoding models to probe the neural substrates responsive to various features by leveraging the multivariate temporal response function (mTRF) [30]. Specifically, the mTRF is a linear model that incorporates time lags (*τ*) between features (*X*) and neural activities (*y*) (Fig. 3B). It can be formulated in a convolutional form:

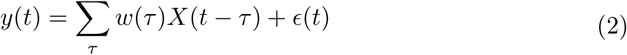

where *w* represents the weights (also known as the mTRF filter) to be estimated, and *ɛ* is the additive noise.

Both neural signals and features were z-scored prior to model training. Given the relatively large parameter spaces (number of lags × feature dimensions), we applied L2-norm regularization during parametric estimation to mitigate the potential over-fitting problem. The encoding model was implemented using four-fold cross-validation (CV), where the data were split into four groups - three groups used for training and the withheld one for testing. Model performance was assessed using Pearson’s correlation (*r* value) between predicted (*y*^) and real signal (*y*). The ridge parameter was optimized using a grid search approach to maximize the r-value during training. Final model performance was calculated as the average r-value across all four folds. In addition, we decided to train mTRF models with delays up to one second (i.e., *τ_max_* = 1 s) for two reasons. First, since only trials longer than 1 second were included, using a consistent mTRF window ensures sufficient data for model estimation. Second, a 1-second window adequately captures key neural processes involved in audiovisual processing during natural movie viewing, including early sensory responses (50–200 ms) [73, 89], audiovisual integration (100–300 ms) [65–67], and semantic processing (∼400 ms) [90]. This window choice is supported by our additional mTRF analyses using progressively increasing lag windows (0–600 ms, 0–800 ms, 0–1000 ms, and –100–1100 ms) for both auditory and visual modalities, which showed no systematic changes across window sizes (*r*-values clustered around the diagonal; Fig. S6). Further, we conducted the encoding analysis on all electrodes rather than the active electrodes that survived from the permutation test, avoiding the potential double dipping problem [91]. Specifically, three types of mTRF models were primarily employed:

*1) Auditory and visual mTRF models.* The auditory and visual mTRF models were trained independently. For each modality, low- and high-level features were first concatenated, followed by a principal component analysis (PCA) procedure to reduce dimensionality to enhance computational efficiency (Fig. 3A). To determine the optimal number of PCs, we trained mTRF models using 5 to 50 principal components (PCs) in increments of 5 PCs (incremental PCs test). Then, we compared the average r-values to identify the elbow point. As a result, this procedure revealed that 15 PCs were optimal (Fig. S4A).
*2) Audiovisual assignment mTRF model.* The averaged global modality feature obtained from the behavior experiment (Fig. 5F) was fed into the encoding model.
*3) Subtitle mTRF model.* The dimension of the subtitle BERT features in the foreign language (FL) condition was also relatively large (i.e., 1024 dimensions). Therefore, PCA was also employed here to reduce the feature dimensions. In the end, 50 PCs were selected for the subtitles, explaining 95.01% variance of the feature space.

Furthermore, we evaluated the statistical significance level of the model performances using a permutation test, wherein the neural data was phase-randomized for 1000 iterations. We repeated the above encoding model training procedure for each permutation to generate a null distribution of r-values. Then, the p-value can be derived by calculating the proportion of r-values in the null distribution that exceeded the real r-value. An electrode was considered significant if *p <* 0.05 and survived the false discovery rate (FDR) correction of multiple comparisons.

### Timing analysis of encoding models

To estimate the latencies of the neural responses, we identified the time delays corresponding to the peak weights in the mTRF filters for each electrode (Fig. S4C). The peak timings were first grouped by ROI (Fig. S1B). Then, we applied a bootstrap procedure to assess the response lags for each ROI. Specifically, we estimated the 95% confidence intervals (CI) by sampling the data with replacement and calculated the mean for 10000 iterations.

A permutation test was conducted to determine statistical significance across different ROIs (Fig. 3E). For each comparison, the group labels of each element were randomly shuffled for 1000 iterations, yielding a null distribution of mean differences. Then, the *p*-value was calculated by determining the position of the real difference in the null distribution. The resulting *p*-value was corrected for multiple comparisons using the FDR method.

### Frontal ventral-dorsal gradient analysis

We constructed the AV index and frontal coordinate to quantify the representational distribution of audiovisual information in the prefrontal cortex. Specifically, the AV index (*I_AV_*) was applied to capture the electrode selectivity for auditory or visual modalities, based on r-values obtained from the auditory and visual mTRF models (*r_A_* and *r_V_*):

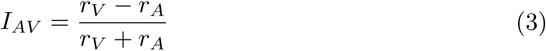

By definition, the range of *I_AV_* is from -1 to 1, where 1 represents the electrode is complete visual tuning and -1 represents complete auditory tuning.

In addition, we established a coordinate system to quantify the ventral-to-dorsal gradient of the frontal cortex. Conventionally, the ventrodorsal organization of the frontal cortex has been characterized based on cytoarchitecture and connectivity, where dlPFC corresponds to Brodmann areas (BA) 46 and 9/46 and vlPFC corresponds to BA 45 and 47/12 [92, 93]. This dichotomy aligns with the division between the inferior frontal gyrus (IFG) and middle frontal gyrus (MFG), demarcated by the inferior frontal sulcus (IFS). Notably, this boundary is not strictly cardinal (i.e., horizontal or vertical), necessitating the use of both *y* and *z* MNI coordinates to accurately capture the ventrodorsal distinction. To this end, we defined a frontal polar coordinate system within the *y*-*z* plane of the MNI space, taking the intersection of precentral sulcus and Sylvian fissure as the origin (MNI coordinate: [-55, 15, -8]; please note that the *x* coordinate was not critical in this analysis). In this system, a smaller radius (*d*, mm) denotes a more ventral position, whereas a larger radius corresponds to a more dorsal location (Fig. 4A, F). This approach also enables us to quantify the modality transition in a fully data-driven manner without imposing rigid anatomical assumptions.

### Encoding model partitioning

We employed a partitioning procedure to obtain the unique effect of a specific feature space [13]. For instance, to quantify the unique effects of the low- and high-level audiovisual features (Fig. 4C), we trained and evaluated models using either low-or high-level features (*M_l_* and *M_h_*), and compared them with models trained using both feature spaces (*M_l∪h_*). Then, the pure contribution of the low-level features was computed as the difference between the combined model and the high-level model (*M_l∪h_* − *M_h_*), and that of high-level features was determined as the discrepancy between the combined model and the low-level model (*M_l∪h_*−*M_l_*). We fitted all mTRF models mentioned above with 15 dimensions using PCA to ensure comparable model complexity.

Similarly, to quantify the unique contributions of audiovisual assignment and features, we trained separate models using either audiovisual assignment (*M_a_*) or audiovisual features (*M_f_*), as well as a combined model incorporating both (*M_a∪f_*). The unique effect of the modality assignment was calculated as the difference between the combined model and the audiovisual features model (*M_a∪f_* − *M_f_*), while the unique contribution of audiovisual features was determined conversely (*M_a∪f_* − *M_a_*; Fig. S10A).

In addition, to determine the influences of the subtitles, we built a full model that combined both the visual and subtitle features. Then, the unique subtitle effect was obtained by calculating the decrease in *r*-values when subtitle features were removed from the full model (Fig. 4D-E; only for the Foreign Language condition).

## Data and Code availability

The dataset of the current study will be made available from the authors upon request and documentation is provided that the data will be strictly used for research purposes and will comply with the terms of our study IRB. The code is available upon publication at https://github.com/flinkerlab/.

## Acknowledgements

We thank A. Ferrari, M. Landy, and S. Michelmann for their comments on an early version of the manuscript, A. Morton for sharing the movie license for academic use, and other members in Flinker lab for extensive discussion. This work was supported by National Institutes of Health grants R01NS109367, R01NS115929, and R01DC018805 (A.F.) and National Science Foundationshum IIS-2309057 (A.F.).

## Author Contributions

F.Z., A.K.-G and A.F. conceived and designed the research; P.D. and D.F. provided clinical care, W.D. and P.R. provided clinical care and performed neurosurgery; A.M. and O.D. provided clinical care and critically revised the article; F.Z. and A.F. wrote the paper.

## Declarations

The authors declare no competing interests.

## Supplementary Information

**Fig. S1:**
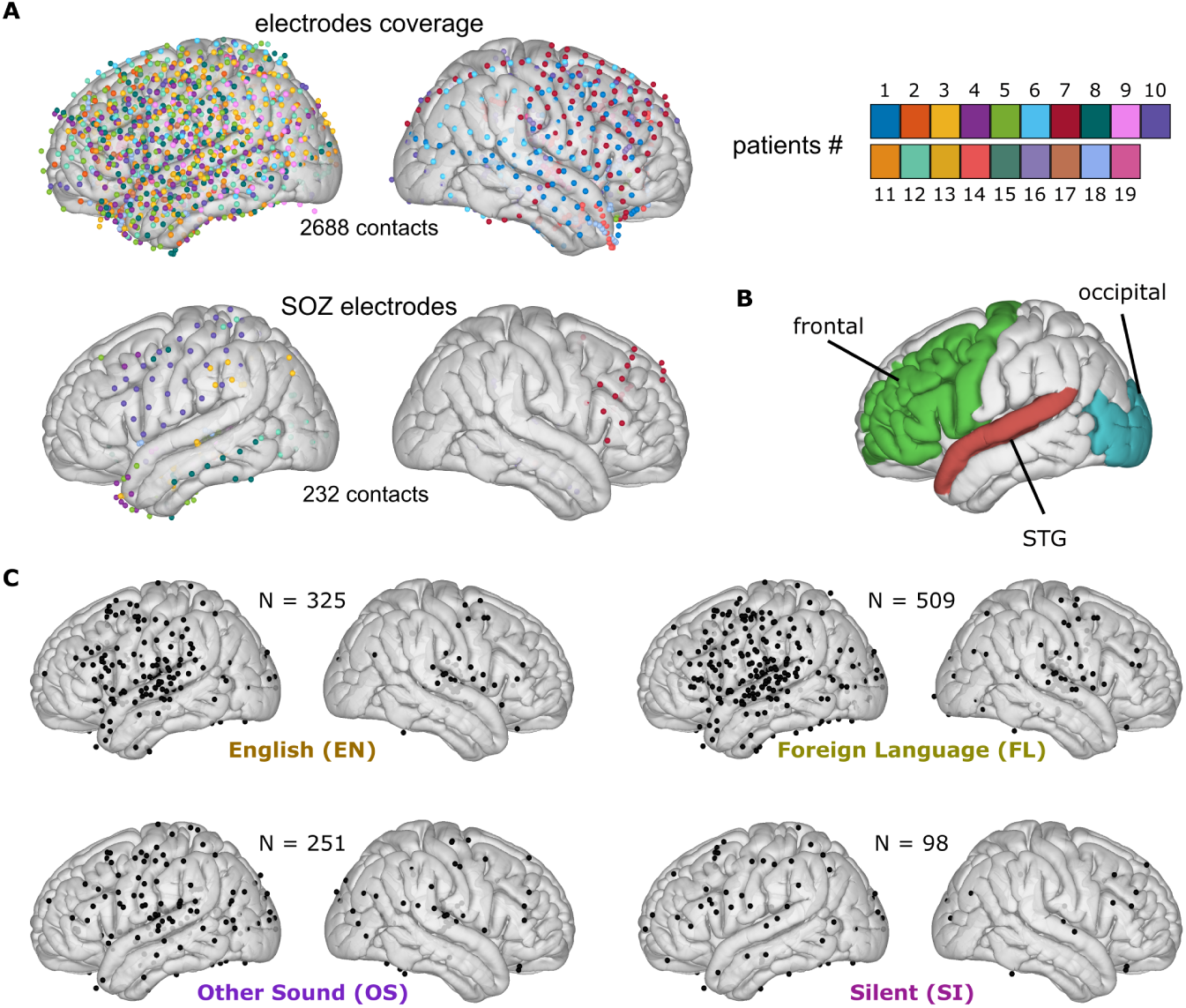
(A) The electrode coverage (upper panel; 2688 contacts in total) and the seizure onset zone (SOZ) electrode distribution (lower panel; 232 contacts in total which were removed from analysis including depth and surface) for all 19 participants involved in the movie viewing task. (B) Three ROIs used in this study, including the superior temporal gyrus (STG), frontal regions (comprising the inferior frontal gyrus, middle frontal gyrus, and pre-central gyrus), and the occipital regions (comprising the lateral occipital gyri, lingual gyrus, and cuneus gyrus). (C) The brain maps of the active electrodes for all four conditions that survived the permutation test (Fig. S2; see Methods: Active electrodes selection).

**Fig. S2:**
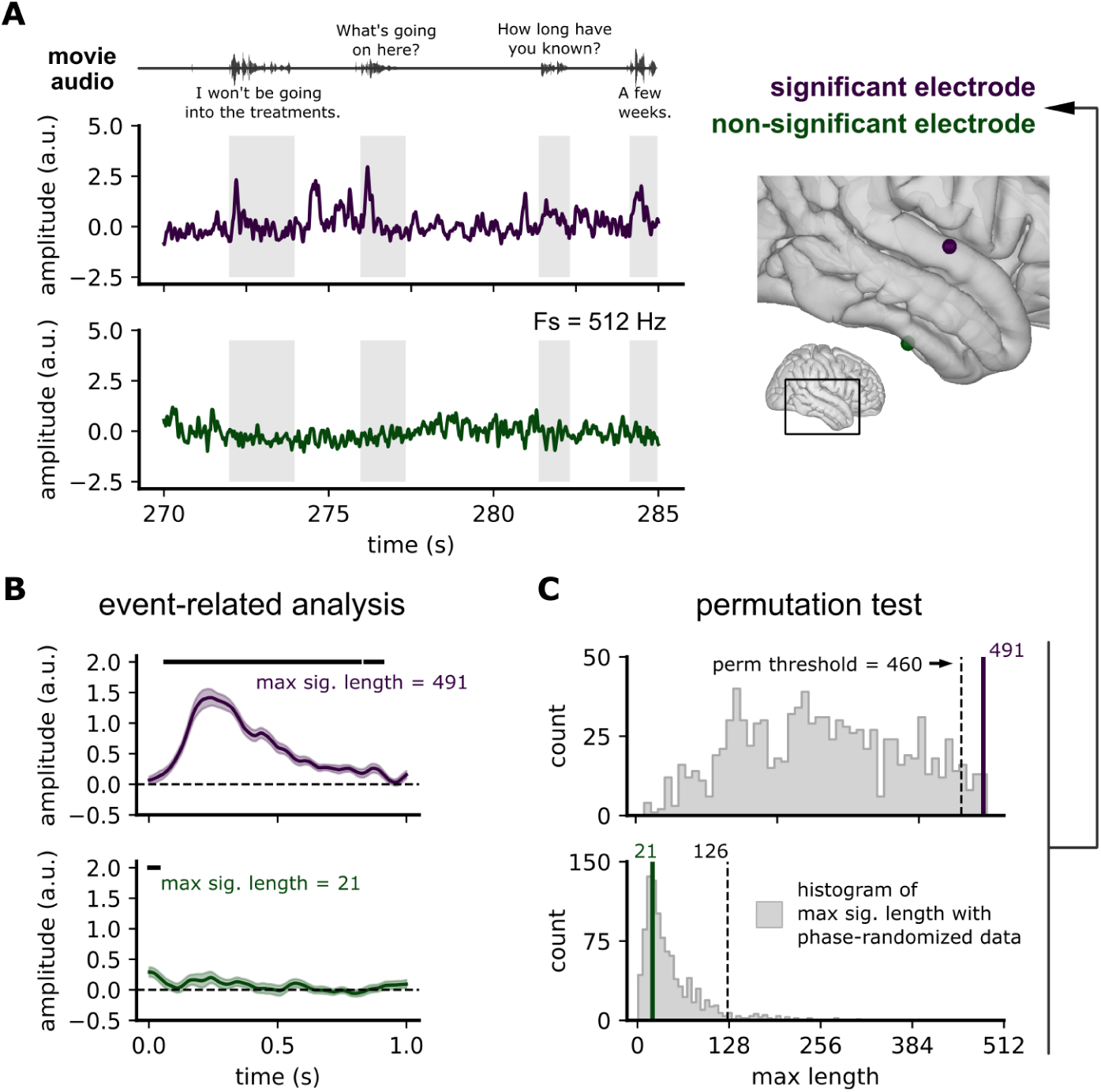
Permutation procedure for selecting the active electrodes. (A) Representative signals from two electrodes (right panel): one located in the STG (purple) and the other located in the ITG (green). The STG electrode exhibited responses to auditory events (left panel), whereas the ITG electrode showed no response. (B) Event-related analysis. The neural activity during the events was aligned to the onset and averaged across time (the temporal window was set as 1 seconds). Then, the statistical significance was determined for each time point using a two-sided *t* -test with *p <* 0.05. The maximum length of the consecutive significant time points (i.e., maximum significant length (*L_max_*)) is recorded. (C) Permutation test. We repeated the event-related analysis on the phase-randomized signals for 1000 times, and computed the maximum significant length for each permutation (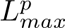). This procedure generated a null distribution of the 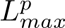. The real length *L_max_* was then compared against this null distribution, with significance determined at the top 5th percentile threshold (FDR corrected).

**Fig. S3:**
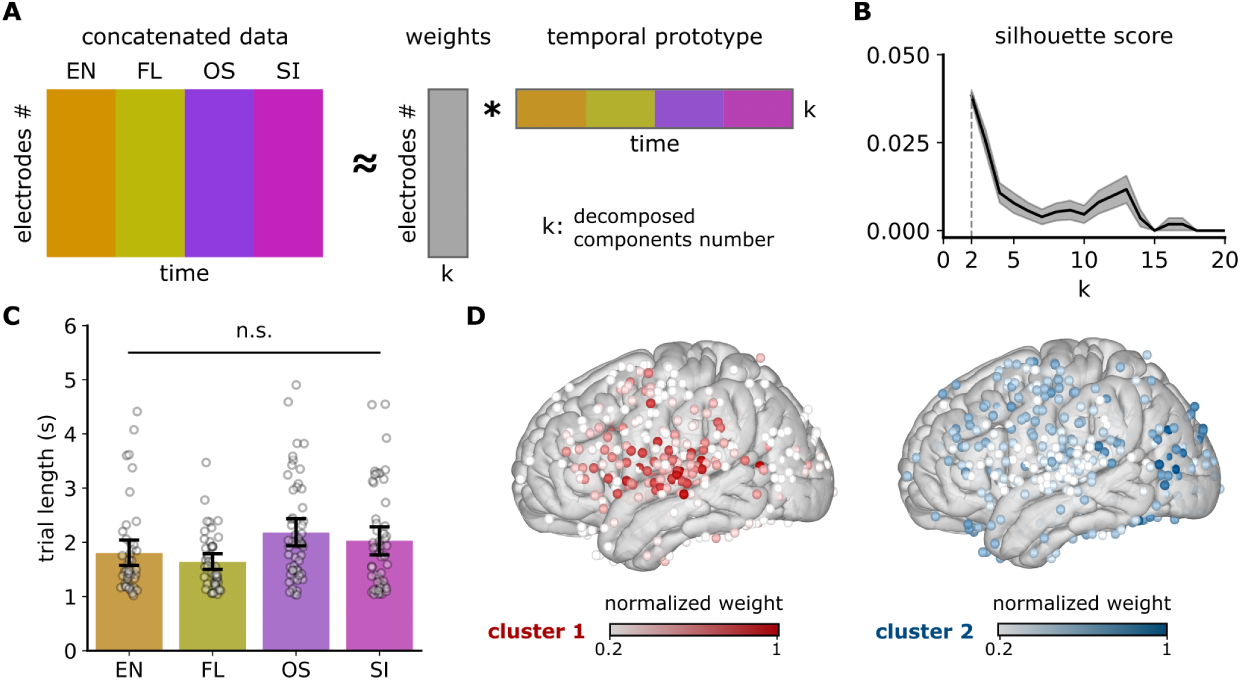
Unsupervised functional clustering. (A)The data for all four conditions were concatenated and then decomposed into a weight matrix and a temporal prototype matrix (see Methods: Functional clustering analysis). (B) The result of silhouette method used to determine the optimal number of clusters (k). Higher scores indicate better clustering performance. The shaded area around the curve represents the standard error (SE). (C) Bar plots of trial lengths across four conditions. Each dot represents an individual trial. (D) The brain maps for the two NMF clusters separately.

**Fig. S4:**
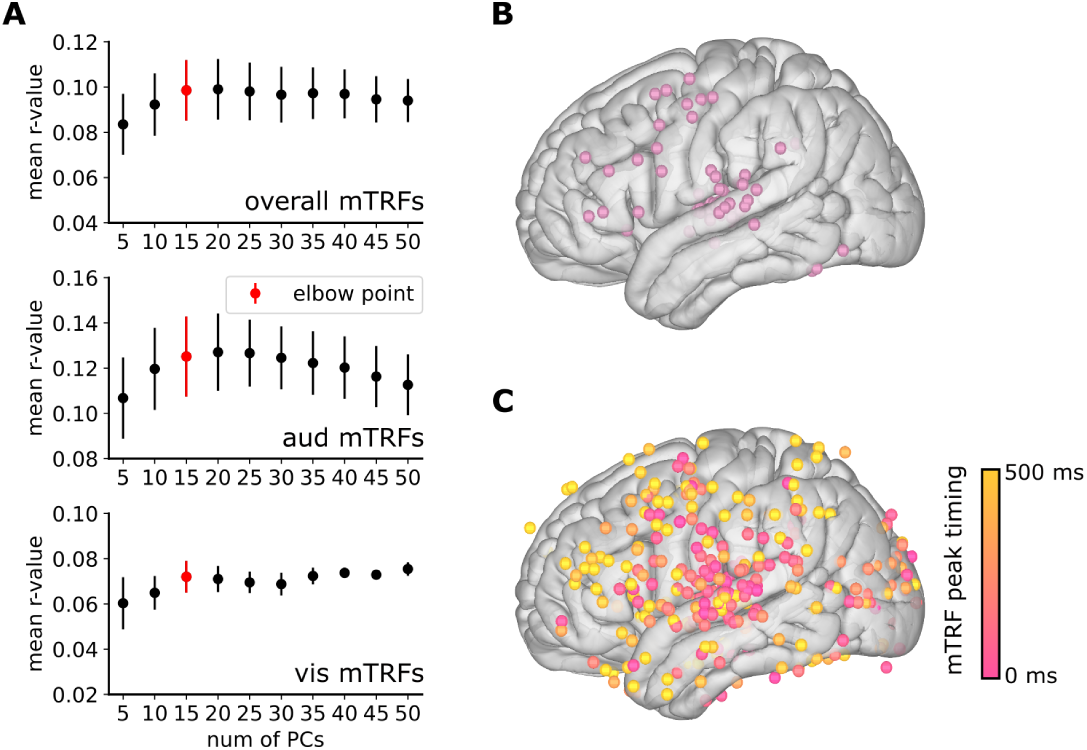
(A) Results of the incremental principal components (PCs) test across all the mTRF models (upper panel), auditory mTRF models (middle panel), and visual mTRF models (lower panel). The red error bars indicate the elbow points identified using kneed Python toolbox, which employs a rotation-based algorithm to detect the maximum curvature [94]. (B) Brain map showing significant electrodes for both audio and visual mTRF models. (C) Timing of peak weights across electrodes that showed significant responses in the auditory and visual mTRF models.

**Fig. S5:**
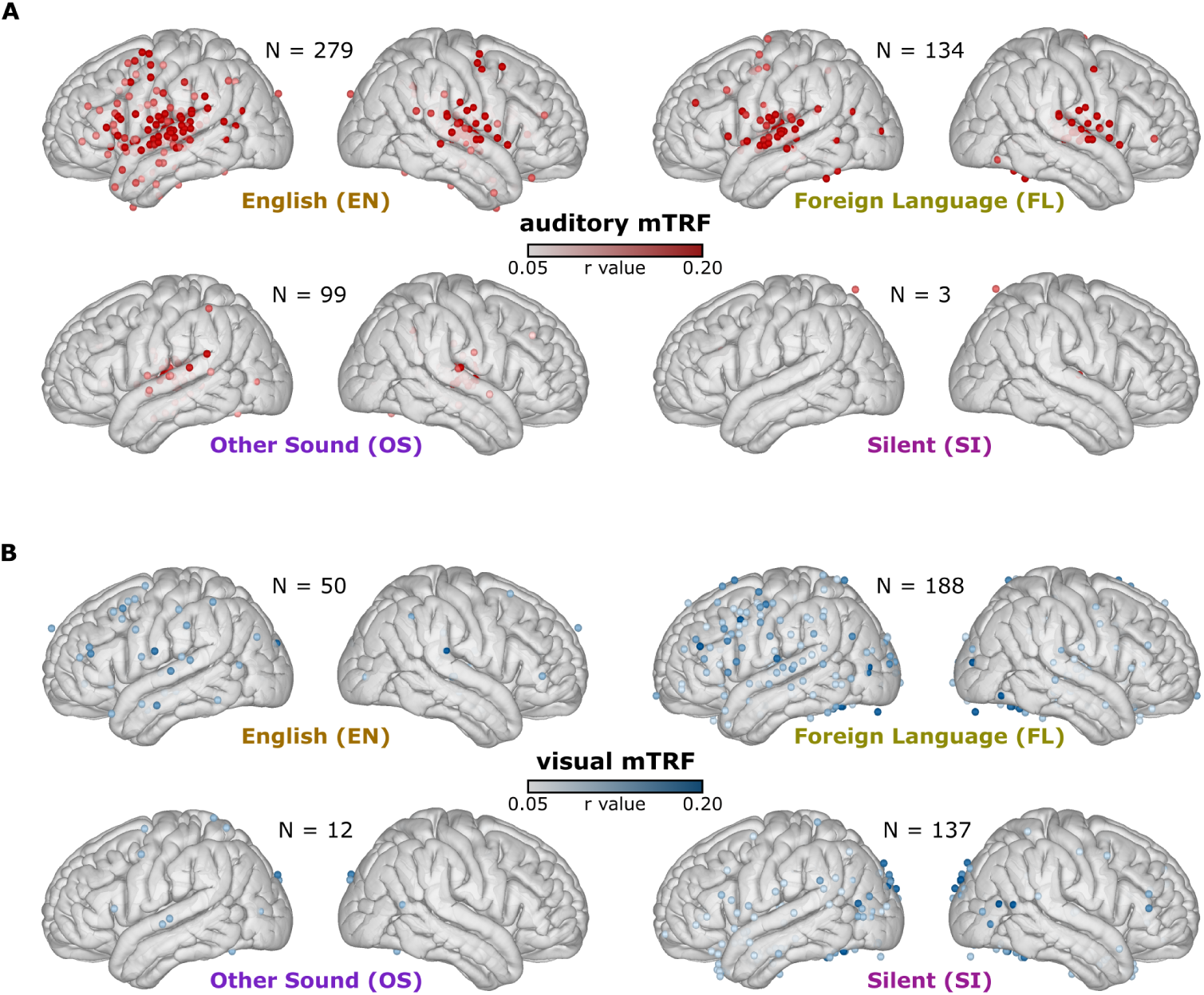
Results of encoding models for auditory (A) and visual (B) information in four conditions. N represents the number of significant electrodes.

**Fig. S6:**
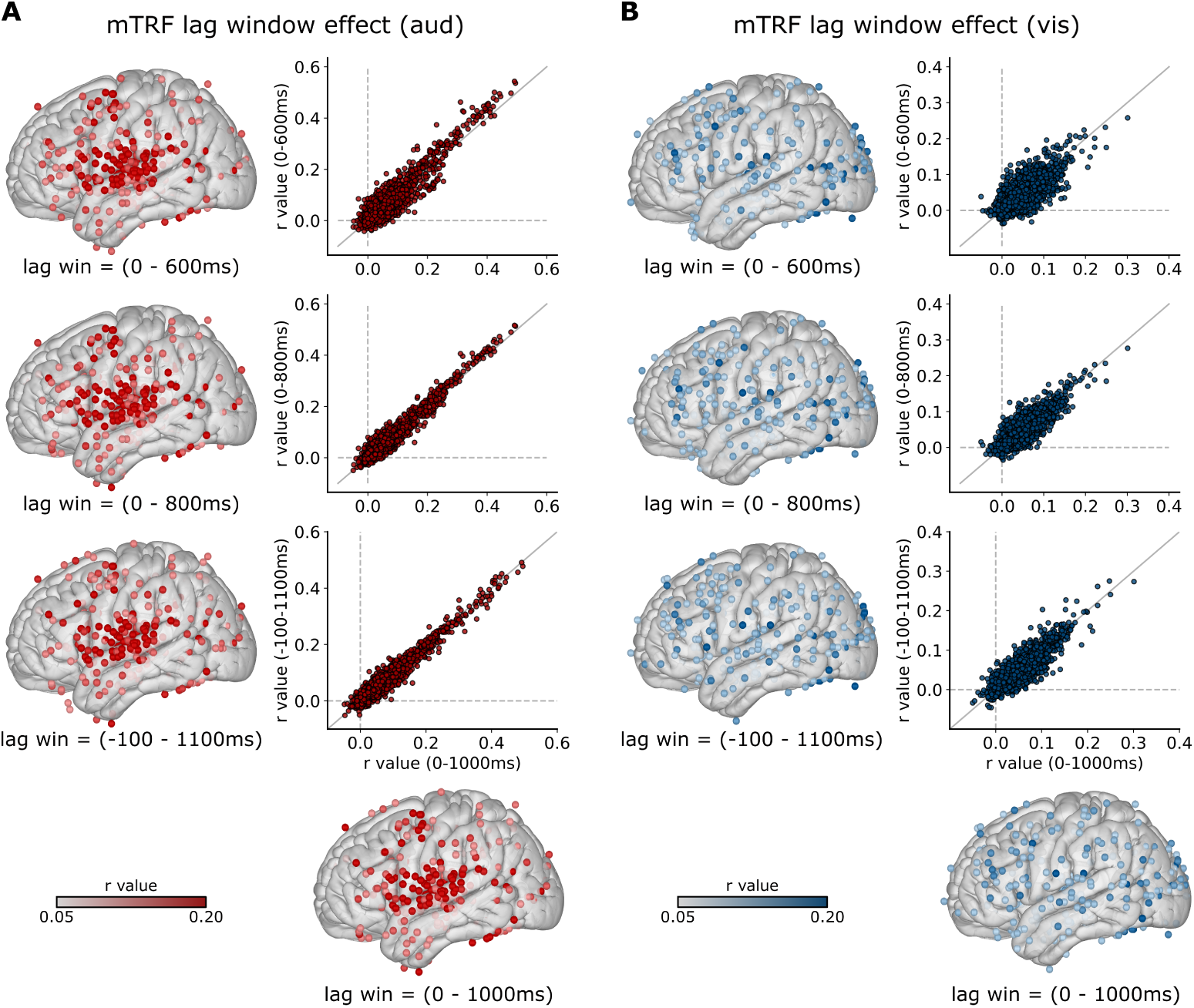
Brain maps and *r*-value comparisons across increasing lag windows (0–600 ms, 0–800 ms, 0–1000 ms, and –100–1100 ms) for both auditory (A) and visual (B) mTRF models. No permutation test or multiple correction were applied; a hard threshold of *r >* 0.1 was used for visualization.

**Fig. S7:**
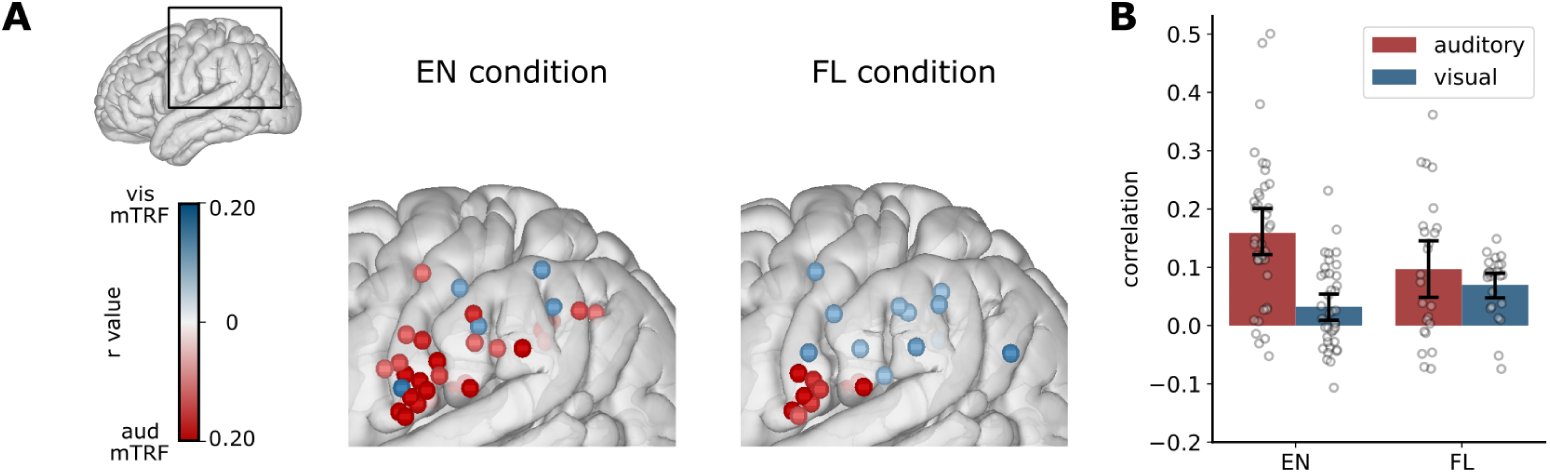
(A) The mTRF correlation maps in the parietal regions (including the post-central gyrus, inferior parietal lobe, and supramarginal gyrus) for the English (EN) and Foreign Language (FL) conditions. (B) Bar plots of the auditory and visual mTRF models in the parietal cortex for the EN and FL conditions. Error bars represent 95% confidence interval (CI).

**Fig. S8:**
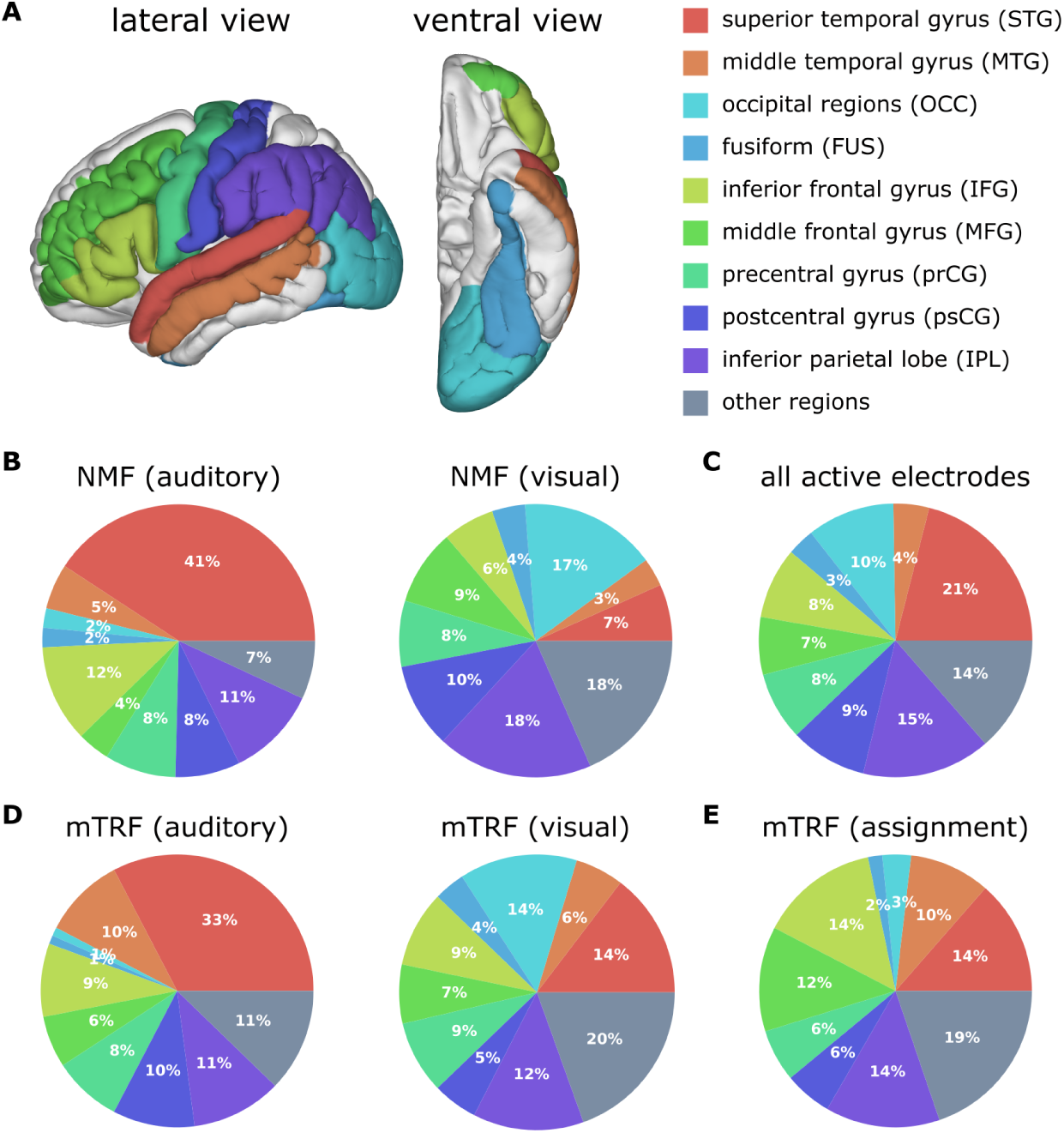
Anatomical distributions for the NMF, mTRF and active electrodes. (A) Nine ROIs illustrated on the brain. (B) Pie charts showing the anatomical distribution of electrodes from the auditory and visual functional clusters (NMF approach). (C) Pie charts showing the anatomical distribution of the active electrodes. (D) Pie charts depicting the anatomical distribution of significant electrodes from the auditory and visual encoding models (mTRF approach). (E) Pie chart showing the anatomical distribution of significant electrodes responsive to the audiovisual assignment (i.e., the global modality). The total number of electrodes within each ROI, and the number (and percentage) of electrodes for figure B-E are shown in Table S2.

**Fig. S9:**
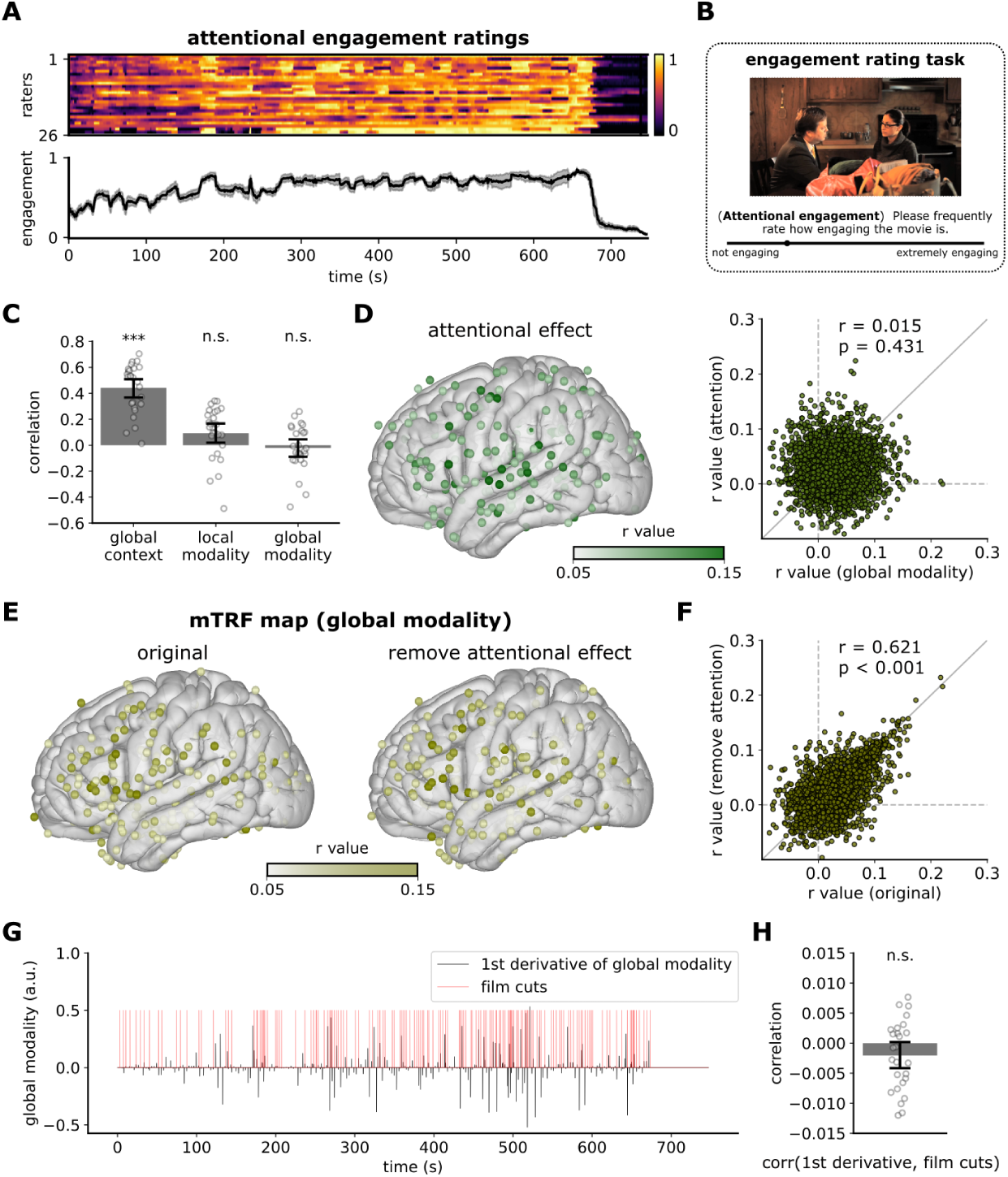
(A) The attentional engagement ratings from all participants over time (upper panel) and the average time series across participants (lower panel). (B) Illustration of the engagement rating task conducted on the Amazon Mechanical Turk platform. (C) Correlations between attentional engagement ratings and global context (one-sample *t* test; *T* (25) = 11.987, *p <* 0.001, *d* = 2.351), local modality (one-sample *t* test; *T* (25) = 2.068, *p* = 0.074, *d* = 0.406), as well as global modality (one-sample *t* test; *T* (25) = −0.608, *p* = 0.549, *d* = 0.119). Each dot represents an individual rater. (D) The mTRF result for the attentional engagement effect. (E) The mTRF results for the original global modality (left) and after removing the attentional effect (right). (F) Comparison of *r* values between the two models across all electrodes. (G) Markers of abrupt changes in the averaged global modality (black; quantified as its first derivative) and the film cuts (red). (H) Correlations between the film cuts and the global modality changes across participants. Each dot represents an individual rater. Error bars in figure C and H represent 95% CI.

**Fig. S10:**
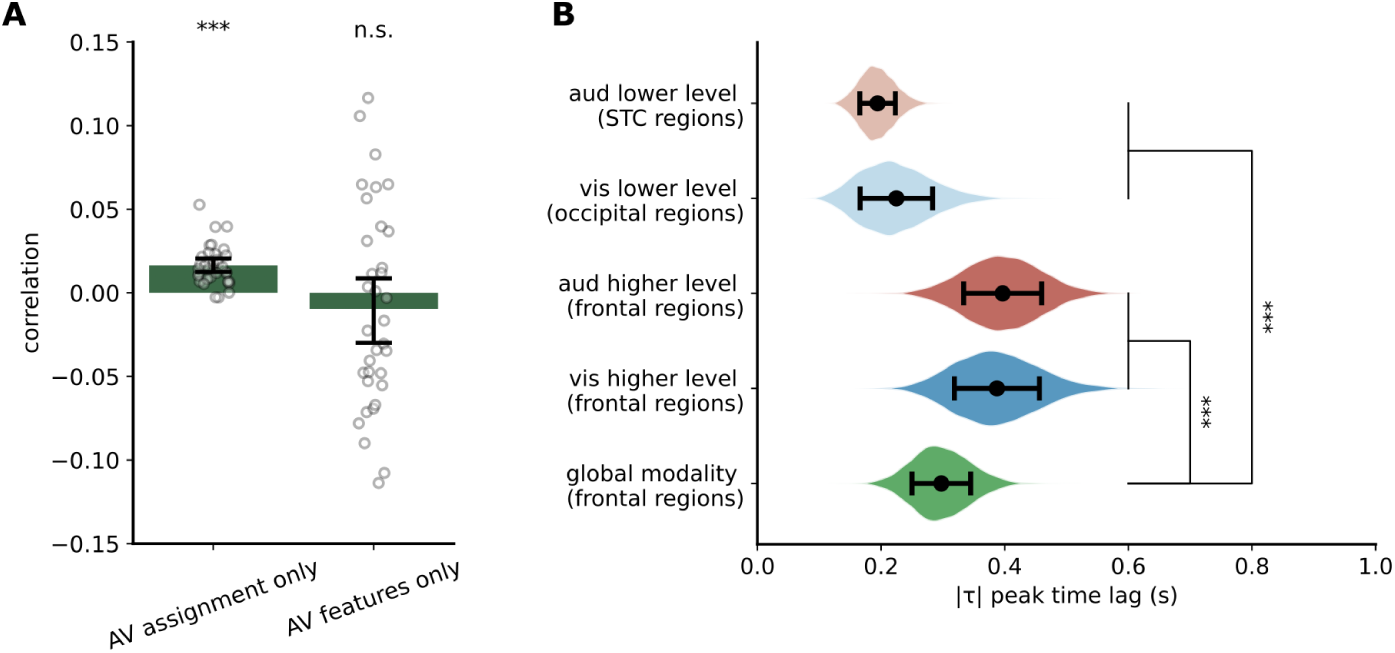
(A) The unique contributions for modality assignment and audiovisual features in frontal electrodes shown in Fig. 6A (see Methods: Encoding model partitioning). A significant effect was observed only for the audiovisual assignment (one sample *t*-test, *t*(33) = 7.968, *p <* 0.001, *d* = 1.366, FDR corrected) but not for the audiovisual features (one sample *t*-test, *t*(33) = −0.927, *p* = 0.361, *d* = 0.159, FDR corrected). The error bars represent 95% CI. (B) Timing analysis of audiovisual (AV) assignment. The timing of AV assignment in the frontal cortex (297 ± 47 ms) was significantly later than the neural representation of AV features in the perceptual regions. A significant difference was also observed between the timing of encoding the AV features and the modality assignment in the frontal regions. The distributions were estimated based on a bootstrap procedure, and the statistical significance between regions was assessed via a permutation test (see Methods: Timing analysis of encoding models). The error bars represent the standard deviation (SD). Significance levels are set as *p <* 0.001 (***), *p <* 0.01 (**), *p <* 0.05 (*), and *p* ≥ 0.05 (n.s.).

**Table S1:** Patient information.

| ID | Coverage | Gender | Handedness | Age at Surgery (y) | Seizure Onset Zone (SOZ) |
| --- | --- | --- | --- | --- | --- |
| 1 | Right | Male | Right | 29 | temporal |
| 2 | Left | Female | Right | 45 | insula/frontal |
| 3 | Left | Male | Right | 24 | parietal |
| 4 | Left | Female | Right | 30 | temporal |
| 5 | Left | Female | Right | 56 | temporal |
| 6 | Bilateral | Male | Right | 26 | temporal |
| 7 | Right | Female | Right | 21 | frontal |
| 8 | Left | Male | Right | 22 | temporal |
| 9 | Left | Male | Right | 44 | temporal |
| 10 | Left | Female | Right | 33 | insula/temporal |
| 11 | Left | Female | Right | 23 | temporal |
| 12 | Left | Female | Right | 18 | occipital |
| 13 | Left | Female | Right | 61 | temporal |
| 14 | Bilateral | Female | Right | 33 | temporal |
| 15 | Left | Female | Right | 27 | temporal/frontal |
| 16 | Right | Male | Right | 28 | insula/temporal/frontal |
| 17 | Left | Male | Right | 26 | parietal |
| 18 | Bilateral | Female | Right | 33 | insula/temporal |
| 19 | Bilateral | Female | Right | 30 | insula/temporal/frontal |

**Table S2:** Electrode anatomical coverage and significant responses.

| ROI | total | active | NMF (aud) | NMF (vis) | mTRF (aud) | mTRF (vis) | mTRF (assign.) |
| --- | --- | --- | --- | --- | --- | --- | --- |
| STG | 146 | 72 (49.32%) | 53 (36.39%) | 12 (8.22%) | 64 (43.84%) | 36 (24.66%) | 24 (16.44%) |
| MTG | 136 | 20 (14.71%) | 7 (5.15%) | 6 (4.41%) | 19 (13.97%) | 14 (10.29%) | 17 (12.50%) |
| OCC | 59 | 32 (54.24%) | 3 (5.08%) | 29 (49.15%) | 2 (3.39%) | 34 (57.63%) | 6 (10.17%) |
| FUS | 22 | 10 (45.45%) | 3 (13.64%) | 7 (31.82%) | 2 (9.09%) | 9 (40.91%) | 3 (13.64%) |
| IFG | 90 | 29 (32.22%) | 15 (16.67%) | 11 (12.22%) | 17 (18.89%) | 22 (24.44%) | 25 (27.78%) |
| MFG | 145 | 24 (16.55%) | 5 (3.45%) | 16 (11.03%) | 12 (8.28%) | 17 (11.72%) | 22 (15.17%) |
| prCG | 101 | 28 (27.72%) | 11 (10.89%) | 14 (13.86%) | 16 (15.84%) | 21 (20.79%) | 11 (10.89%) |
| psCG | 106 | 36 (33.96%) | 10 (9.43%) | 18 (16.98%) | 19 (17.92%) | 13 (12.26%) | 10 (9.43%) |
| IPL | 208 | 51 (24.52%) | 14 (6.73%) | 33 (15.87%) | 21 (10.10%) | 32 (15.38%) | 24 (11.54%) |
| Other regions | 293 | 49 (16.72%) | 9 (3.07%) | 33 (11.26%) | 24 (8.19%) | 48 (16.38%) | 35 (11.95%) |
| Other (WM & UNK) | 1150 | 299 (26%) | 35 (3.04%) | 19 (1.65%) | 123 (10.70%) | 44 (3.83%) | 129 (11.22%) |

## Notes

### Competing Interest Statement

The authors have declared no competing interest.

### Summary of Updates

Figure 2 and 4 are revised. Supplementary figure 1, 3, 4, 6, 7, 9 are also updated. Additional control analyses have been described and discussed in the main text.

